# The (p)ppGpp-binding GTPase Era promotes rRNA processing and cold shock survival in *Staphylococcus aureus*

**DOI:** 10.1101/477745

**Authors:** Alison Wood, Sophie E. Irving, Daniel J. Bennison, Rebecca M. Corrigan

## Abstract

Ribosome assembly cofactors are widely conserved across all domains of life. One such group, the ribosome-associated GTPases (RA-GTPase), act as molecular switches to coordinate ribosome assembly. We previously identified the *Staphylococcus aureus* RA-GTPase Era as a target for the stringent response alarmone (p)ppGpp, although the function of Era in ribosome assembly is unclear. Era is highly conserved throughout the bacterial kingdom and is essential in many species. Here we show that Era is not essential in *S. aureus* but is important for 30S ribosomal subunit assembly. Protein interaction studies reveal that Era interacts with the endonuclease YbeY and the DEAD-box RNA helicase CshA. We determine that both Era and CshA are required for growth at suboptimal temperatures, virulence and 16S rRNA processing. Era and CshA also form direct interactions with the (p)ppGpp synthetase RelSA, with RelSA positively impacting the GTPase activity of Era. We propose that Era acts to direct enzymes involved in rRNA maturation and ribosome subunit assembly to their site of action, an activity that is regulated by components of the stringent response.

**Author summary:** The bacterial ribosome is an essential cellular component and as such is the target for a number of currently used antimicrobials. Correct assembly of this complex macromolecule requires a number of accessory enzymes, the functions of which are poorly characterised. Here we examine the function of Era, a GTPase enzyme involved in 30S ribosomal subunit biogenesis in the important human pathogen *S. aureus*. We uncover that Era is not an essential enzyme in *S. aureus*, as it is in many other species, but is important for correct ribosome assembly. In a bid to uncover the cellular function of this enzyme, we identify a number of protein interaction partners with roles in ribosomal RNA maturation and degradation, supporting the idea that Era acts as an intermediary protein facilitating ribosomal biogenesis. We also uncover a link between Era and the (p)ppGpp synthetase RelSA, revealing an additional level of control of rRNA levels by the stringent response. With this study we elaborate on the functions of GTPases in ribosomal assembly, processes that are controlled at multiple points by the stringent response.

## Introduction

Ribosomes are macromolecular machines responsible for the synthesis of proteins in all living cells. In bacteria these complexes consist of two subunits, with the large 50S subunit containing 33 large proteins (L1-36) and two rRNAs, and the 30S small subunit containing 21 small proteins (S1-21) and the 16S rRNA. As such, assembly of the ribosome is a tightly regulated process, with correct maturation requiring the help of assembly cofactors, one class of which are the P-loop ribosome-associated GTPases (RA-GTPase). These enzymes are widely conserved across all domains of life and act as molecular switches, cycling between inactive GDP-bound, and active, effector-binding GTP-bound states. Within the P-loop GTPase class lies the Era family (*<u>E</u>scherichia coli* <u>Ra</u>s-like protein), which is characterised by the presence of a distinct derivative of a KH domain [1]. Era, the protein after which this family is named, is highly conserved throughout the bacterial kingdom, although is missing in *Chlamydia* and mycobacterial species [1]. This GTPase is essential in *E. coli* [2–5], *Salmonella* Typhimurium [6] and in strains of *Bacillus subtilis* [7, 8]. The first indication that Era is involved in ribosomal biogenesis came from the ability of the 16S rRNA methyltransferase KsgA to suppress a cold-sensitive phenotype of an Era E200K mutation [9]. While important for ribosomal maturation, additional phenotypic defects associated with depleted cellular levels of Era include cell cycle control and chromosome segregation, as well as carbon and nitrogen metabolism [10–12].

Era is composed of two domains, an N-terminal GTPase domain and a C-terminal RNA-binding K-homology domain [13]. A cryo-electron micrograph structure of Era in complex with the 30S subunit of the ribosome reveals that Era binds into the same pocket as small subunit protein S1 [14]. In the absence of S1, Era interacts with proteins S2, S7, S11 and S18, as well as with a number of helices of the 16S rRNA. In addition, Era interacts with h45 and nearby residues 1530-1539 (GAUCACCUCC) in the 3’ minor domain of the 16S rRNA via its CTD region [14, 15]. These residues include the anti-Shine Dalgarno sequence, critical for the formation of the 30S preinitiation complex. Including Era in *in vitro* reconstitutions of the 30S ribosome promotes the incorporation of several late-stage ribosomal proteins for the RNA [16, 17]. Consequently, it has been proposed that Era functions as a checkpoint protein, and that by binding to the 16S rRNA the formation of the initiation complex is prevented until the appropriate time [14]. In addition to interacting with ribosomal proteins, Era has also been reported to interact with a number of proteins involved in 16S rRNA maturation. One of these, YbeY, is an endonuclease required in *E. coli* for the maturation of the 3’ end of the 16S rRNA [18, 19]. It is proposed that the binding of YbeY to Era and S11 guide the endonuclease to its site of action [18].

The stringent response is a bacterial signalling system used by bacteria to cope with a variety of environmental stresses, the best characterised of which is nutrient deprivation. The opportunistic pathogen *Staphylococcus aureus* contains three enzymes, RelSA, RelP and RelQ, which upon sensing a stress, synthesise the nucleotides guanosine tetra- and pentaphosphate ((p)ppGpp) [20, 21]. Once produced, this alarmone controls cellular responses to aid survival. Our previous work identified Era, as well as three other GTPase enzymes from *S. aureus*, as target proteins for (p)ppGpp and demonstrated that the production of (p)ppGpp has a negative impact on mature 70S assembly [22]. Here, we examine the role of Era as an enzyme required for ribosome biogenesis. Unlike in *E. coli*, this GTPase is not essential for the growth of *S. aureus*, however mutant cells are defective in 30S subunit maturation. We identify the *S. aureus* endonuclease YbeY and the DEAD-box RNA helicase CshA as interaction partners for Era, and show that both Era and CshA are crucial for rRNA homeostasis and growth at low temperatures. We additionally demonstrate that both Era and CshA interact with the (p)ppGpp synthetase RelSA, and that cellular rRNA processing is controlled in a stringent response-dependent manner. With this, we propose that Era is a protein that facilitates the interactions between a number of rRNA processing and degrading enzymes and the 30S subunit/16S rRNA, and show that under stress conditions the stringent response is important for this process.

## Results

### Era is not essential in *S. aureus* but is required for normal growth and ribosome assembly

Previous reports show that *era* is an essential gene in multiple bacterial species [3, 6]. In agreement with this, transposon insertions in this gene have not been identified in a number of published *S. aureus* transposon libraries [23–25]. Closer inspection of the transposon insertion hits within the *era* operon obtained in the Nebraska *S. aureus* mutant library [23] reveals insertions in *ybeZ*, a gene of unknown function, and at the 3’ end of the diacylglycerol kinase *dgkA* but not within *ybeY, cdd, era* or *recO*, suggestive of an additional promoter upstream of *era* and *cdd* (Fig 1a). To rule out polar effects and examine whether *era* in isolation is essential, we replaced the entire coding sequence with the Tet-encoding *tetAM* open reading frame in the presence of the anhydrotetracycline (Atet)-inducible covering plasmid pCN55iTET-era. Following deletion of the chromosomal copy of *era*, we attempted to phage transduce this mutation into *S. aureus* strains containing the empty vector pCN55iTET or the complementing plasmid pCN55iTET-era. Transduction efficiencies were similar with both recipients, albeit smaller colony sizes were observed in the absence of *era*, indicating that this gene is not essential in *S. aureus*. We next transduced the *era* deletion into the community-associated methicillin resistant *S. aureus* strain LAC* to rule out secondary mutations and used this strain for further studies. Analysis of growth rates revealed that an *era* mutant strain, while viable, does have a significant growth defect, which can be complemented by the expression of Era from pCN55iTET-era (Fig 1b).

**Fig 1.**
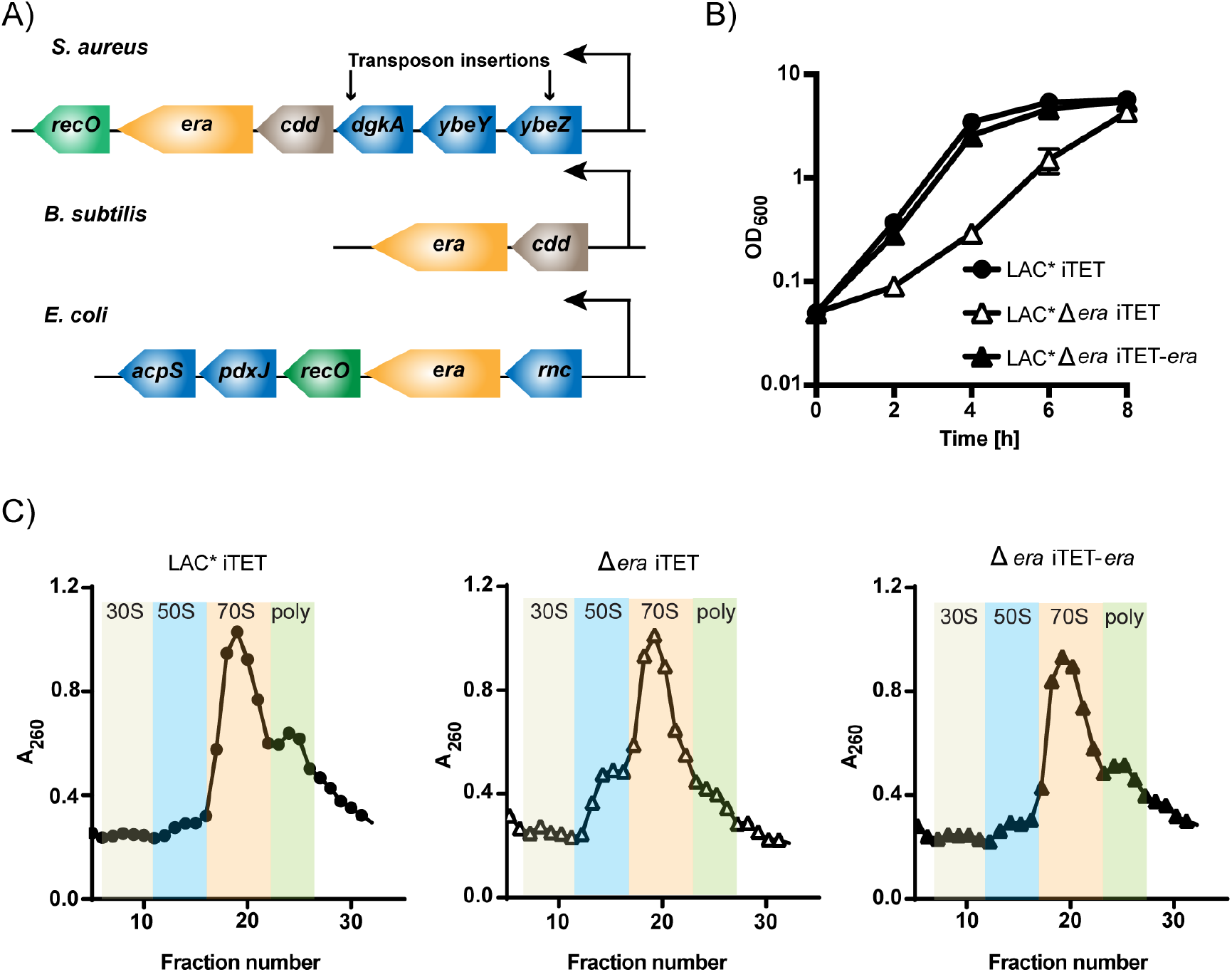
Role of Era in bacterial growth and ribosome assembly. A) Schematic representation of the era-containing operons from *S. aureus, B. subtilis* and *E. coli*. The *era* operon in *S. aureus* consists of: *ybeZ* – encoding for a protein of unknown function; *ybeY* – a 16S rRNA endoribonuclease; *dgkA* - a diacylglycerol kinase involved in lipid metabolism;*cdd* - a cytidine deaminase involved pyrimidine metabolism; *era*, and *recO* encoding for a DNA repair protein. Arrows indicate the site of transposon insertions in the *S. aureus* Nebraska Transposon library. B) Growth of *S. aureus* strains LAC* iTET, LAC* Δ*era* iTET and LAC* Δ*era* iTET-era. Overnight cultures were diluted to an OD_600_ of 0.05 and grown in the presence of 100 ng/ml Atet for 8 h. Growth curves were performed in triplicate (N=3), with averages and standard deviations shown. C) Ribosome profiles from LAC* iTET, LAC* Δ*era* iTET and LAC* Δ*era* iTET-era. Normalised extracts from each strain were layered onto 10-50% sucrose gradients. Gradients were fractionated and analysed for RNA content at an absorbance of 260 nm. 30S, 50S, 70S and polysomes-containing fractions are indicated. Experiments were performed in triplicate (N=3) with one representative graph shown.

A role for Era as a ribosomal subunit assembly cofactor has been reported in both *E. coli* and *B. subtilis*, where depletion leads to a decrease in 70S ribosomes and an accumulation of individual 50S and 30S subunits [26, 27]. We reasoned that the growth defect observed in *S. aureus* may be due to defects in ribosome assembly. To investigate this in the context of a complete *era* deletion, we analysed the cellular ribosomal content of wildtype, mutant and complemented strains by sucrose density centrifugation. This revealed that the *era* mutant strain contained fewer polysomes with a concurrent build-up of 50S subunits, a defect that was reversed in the presence of the complementing plasmid (Fig 1c). As cryo-electron microscopy has shown Era interacting with the 30S subunit [14], an excess of free 50S subunits may be, in and of itself, an indication that there is a defect in small subunit biogenesis, leading to a build-up in free 50S. While the peak for polysomes disappeared, the levels of 70S ribosomes remained similar between the wildtype and mutant, however it is not known if these 70S ribosomes are fully functional. We reasoned that this defect in ribosome assembly might make the *era* mutant strain more susceptible to ribosome-targeting antibiotics. In agreement with this, we observed that the minimum inhibitory concentration (MIC) for the Δ*era* strain decreased 2-4-fold when exposed to the 30S-targeting antibiotic spectinomycin (Fig S1a). However, the MIC was unaffected by the 50S targeting antibiotic chloramphenicol (Fig S1b). Together this indicates that while Era is not an essential protein in *S. aureus*, it is important for optimal growth and ribosome maturation.

### Uncovering Era protein interaction dynamics in a native background

In *S. aureus*, Era is encoded in an operon with five other genes, one of which encodes YbeY (Fig 1a). YbeY is an endoribonuclease implicated in the maturation of the 3’ terminus of 16S rRNA in *E. coli*, as well as a quality control checkpoint protein that together with RNase R is involved in eliminating defective 70S ribosomes [19, 28]. Era interacts with YbeY in *E. coli* [18], although these two genes are not encoded in the same operon in that organism (Fig 1a). To determine whether Era from *S. aureus* interacts with YbeY, and/or any other proteins encoded in the *S. aureus era* operon, we first used a bacterial two-hybrid approach, heterologously expressing Era in combination with YbeZ, YbeY, DgkA, Cdd or RecO in *E. coli*. Using this approach, we observed that Era potentially interacts with YbeY and YbeZ (Fig 2a).

**Fig 2.**
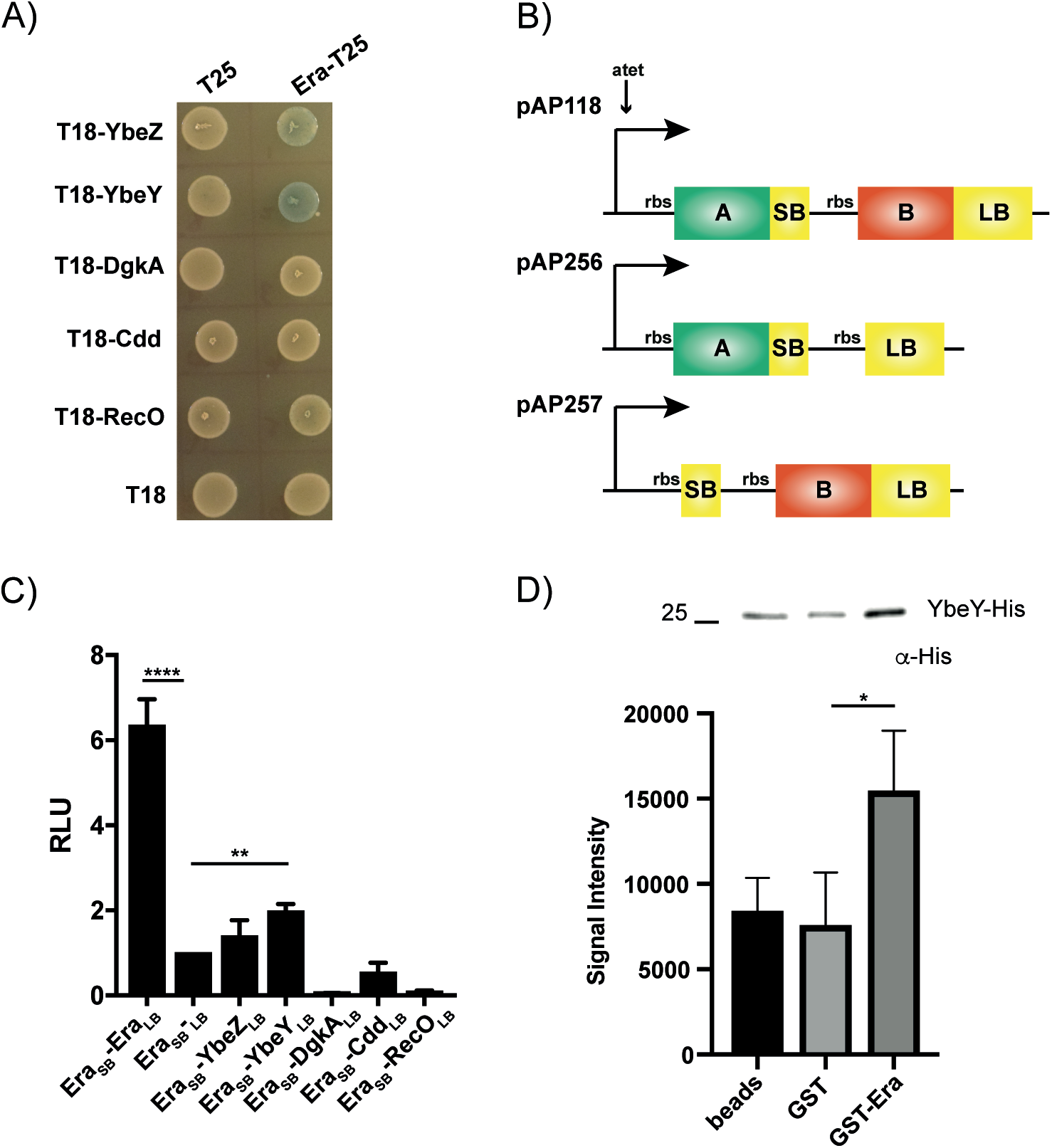
Era interacts with other proteins encoded in the same operon. A) Bacterial two-hybrid showing a positive interaction between Era-T25 and both T18-YbeZ and YbeY. T25 and T18-encoding empty vectors are used as negative controls. N=3, with one representative image shown. B) Schematic representation of the luciferase vectors used for native protein-protein interaction studies. pAP118 contains an Atet-inducible promoter upstream of two genes – SB encodes for the small bit of the luciferase protein, while LB encodes for the large bit. When genes encoding for interacting proteins (A & B) are translationally fused to SB and LB they associate to produce luciferase. pAP256 and pAP257 are control vectors. C) Split luciferase assay demonstrating a significant interaction between Era-Era and Era-YbeY in *S. aureus*. The negative is pAP256-*era*_SB-LB_, which has Era fused to the SB but nothing fused to LB. The negative is normalised to a relative luminescence unit (RLU) of 1. The average values and standard deviations of triplicate experiments are plotted (N=3). Statistical analysis was performed using a one-way ANOVA, followed by Dunnett’s multiple comparisons test (** *P* < 0.005, **** *P* < 0.0001). D) Top: affinity pulldown assay using GST or GST-tagged Era coupled to glutathione beads. Beads alone, GST or GST-Era coupled beads were incubated with His-tagged YbeY. After washing, bound YbeY-His was detected using HRP-conjugated anti-His antibodies. N=3 with one representative image shown. Below: the mean signal intensities and standard deviations from the 3 repeats are plotted. Statistical analysis was performed using a one-way ANOVA, followed by Dunnett’s multiple comparisons test (* *P* < 0.05).

In order to look at these interactions natively in an *S. aureus* background, we adapted a split-luciferase system recently developed to analyse protein-protein interactions in *Clostridium difficile* for use in *S. aureus* (Fig 2b) [29]. The *era* gene, in combination with itself and each of the operon genes, were cloned into the split-luciferase plasmid pAP118 and introduced into *S. aureus* strain LAC*. Induction of protein expression with Atet revealed a strong positive interaction for Era with itself (Fig 2c), indicating that Era can form dimers. Dimerisation of Era has previously been observed while solving the crystal structure of the *E. coli* protein [30], and so this interaction confirms the functionality of the split-luciferase system in *S. aureus*. In addition, a significant interaction was also observed for Era and YbeY (2 x above the control), although not for YbeZ (1.40 x above the control) (Fig 2c). In agreement with the bacterial two-hybrid results, no interactions occurred between Era and DgkA, Cdd or RecO. A decrease in luminescence below the Era control pAP256-era_SB_-LB occurred when co-expressing Era with DgkA and RecO. The reason for this is unclear but could be due to toxicity upon over-expression of a second copy of these genes from a multi-copy plasmid. To confirm the validity of the Era-YbeY interaction *in vitro*, we performed pulldown assays using glutathione beads coupled to either GST or GST-tagged Era and incubated them with His-tagged YbeY (Fig 2d). Here we observed low-level cross reactivity of YbeY with the gluthatione beads that could not be reduced despite increased washes. However, His-YbeY was pulled down significantly more in the presence of Era (Fig 2d). Altogether the bacterial two-hybrid, split luciferase and affinity pulldown assays confirm an interaction between Era and the endonuclease YbeY from *S. aureus*. With this data we have adapted a luciferase system for confirmation of protein-protein interactions in a native *S. aureus* background.

### Era interacts with the DEAD-box RNA helicase CshA

We wished to shed further light on the role played by Era in the cell and sought to identify unknown interaction partners using a genome-wide bacterial two-hybrid screen. A library of *S. aureus* genomic DNA fragments cloned into pUT18C was screened against pKNT25-era. Of the 17 hits obtained, 14 contained fragments mapping to *cshA*, a gene encoding a DEAD-box RNA helicase. CshA functions in mRNA protection and RNA decay in *S. aureus* [31, 32], while the homologue CsdA from *E. coli* is involved in the cold shock degradosome [33]. To confirm the interaction between Era and CshA, full-length *cshA* was cloned into both pUT18 and pUT18C. Upon co-transformation, interactions occurred between CshA-T18 and both T25-Era and Era-T25 (Fig 3a).

**Fig 3.**
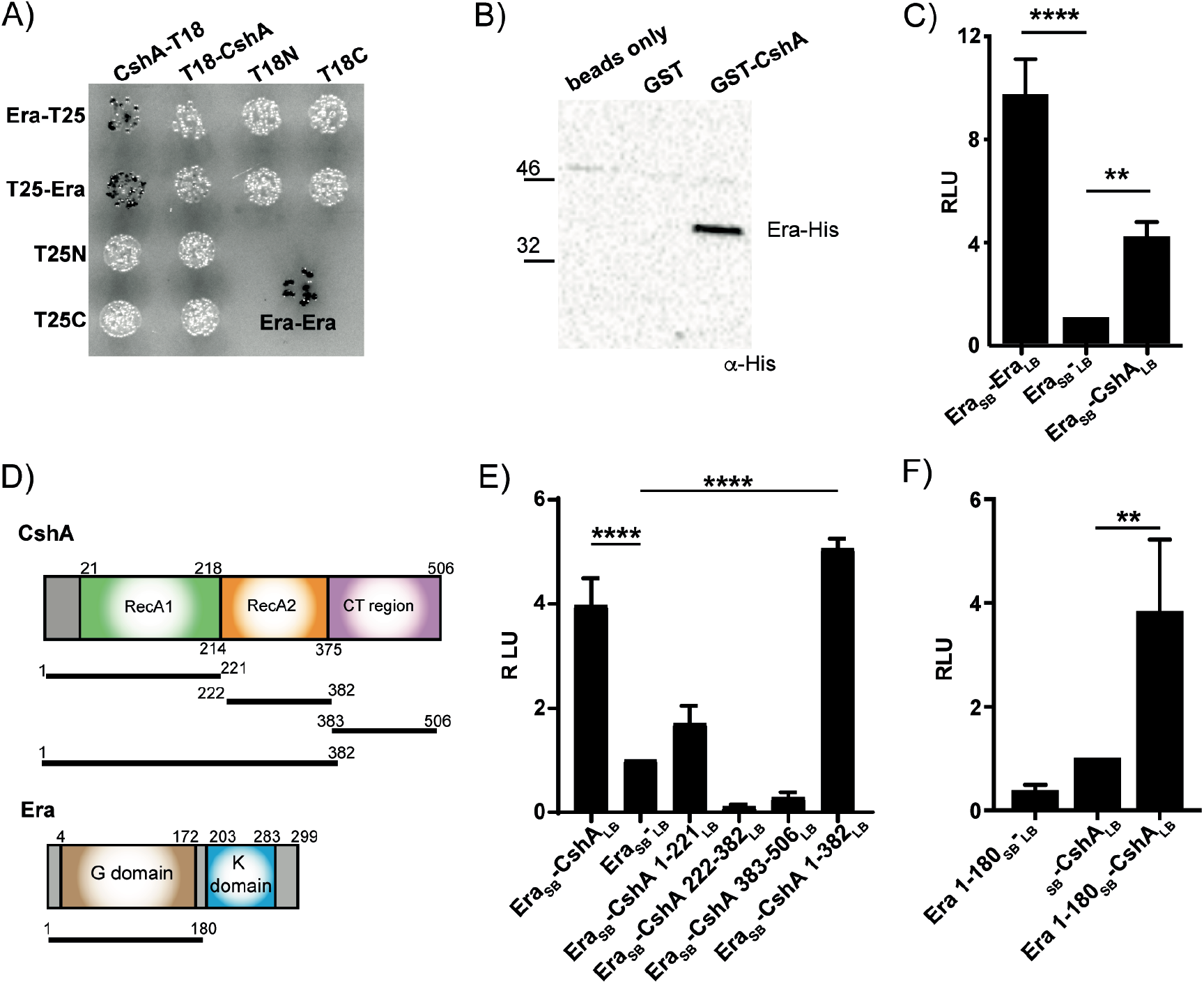
Era interacts with the DEAD-box helicase CshA. A) Bacterial two-hybrid showing an interaction between CshA-T18 and both Era-T25 and T25-Era. Era is known to form a dimer and an Era-Era interaction is used as a positive control. N=3 with one representative image shown. B) Affinity pulldown assay using GST or GST-tagged CshA coupled to glutathione beads. Beads were incubated with His-tagged Era and after washing bound protein was detected using HRP-conjugated anti-His antibodies. N=3 with one representative image shown. C) Split luciferase assay demonstrating an interaction between Era and CshA in *S. aureus*. Dimers resulting from Era fused to both the small (SB) and large bits (LB) of the luciferase gene function as a positive control. The negative, which is normalised to a relative luminescence unit (RLU) of 1, has Era fused to the SB but nothing fused to LB. Cells were grown in the presence of 100 ng/ml Atet at 37°C before being normalised. The average values and standard deviations of triplicate experiments (N=3) are plotted. D) Schematic representation of CshA and Era. The sizes of each domain construct are indicated by the black lines and numbering. E) Split luciferase assay with Era fused to SB and various truncated domain constructs of CshA fused to LB. F) Split luciferase assay with the G domain of Era (1-180) fused to SB and full-length CshA fused to LB. The average values and standard deviations of triplicate experiments (N=3) for all luciferase assays are plotted. All statistical analysis was performed using a one-way ANOVA, followed by Dunnett’s multiple comparisons test (** *P* < 0.005, **** *P* < 0.0001).

To assess this interaction *in vitro*, we performed pulldown assays using glutathione beads coupled to either GST or GST-tagged CshA and incubated them with His-tagged Era, revealing that while His-Era did not interact with the control GST protein, it was pulled down in the presence of CshA (Fig 3b). Finally, to analyse and confirm this interaction in the context of the native host, we used the split luciferase assay (Fig 2b). Assaying for luminescence revealed a 4-fold increase in luciferase activity upon co-expression of Era and CshA (Fig 3c), confirming the identification of a novel Era interaction partner.

### The N-terminal region of CshA is crucial for interacting with Era

CshA is a DEAD-box RNA helicase with an N-terminal helicase core containing two RecA-like domains and a disordered C-terminal (CT) region involved in RNA binding (Fig 3d) [32]. This protein is capable of unwinding both double stranded RNA and RNA-DNA hybrids and is required for both the stabilisation and the degradation of mRNA, the latter of which occurs via interactions of the CT region with components of the RNA degradosome [31, 32].

To determine which regions of both Era and CshA are important for interacting, shorter domain constructs comprising only RecA1 (aa 1-221), RecA2 (aa 222-382), the entire core helicase domain (aa 1-382) or just containing the disordered CT-region (aa 383-506) were cloned, together with Era, into the split luciferase vector pAP118. Interactions with full-length Era were only apparent in the presence of the full core helicase domain (aa 1-382: Fig 3e), indicating that the RNA-binding CT region of CshA is dispensable for this interaction. To determine whether the GTPase domain of Era is required for binding, a shorter construct comprising the N-terminal G domain (aa 1-180) was cloned into pAP118 alongside full-length CshA. Luciferase assays reveal that the G domain is sufficient for this interaction (Fig 3f).

### Interactions between Era and CshA do not affect enzymatic activity

CshA is an RNA helicase, while Era has GTPase activity. To investigate whether the interaction between these two proteins affects the enzymatic activity of either one, we first performed helicase assays. A double stranded RNA oligo was incubated with each protein singly or in combination. While CshA was able to unwind the dsRNA, the addition of Era had no effect on its activity (Fig S2a). In addition, the GTPase activity of Era was unaltered in the presence of either CshA or YbeY (Fig S2b and S2c), indicating that while these proteins do interact, this binding has little effect on their enzymatic functions. As previously proposed, we suggest that these interactions fit with a role for Era as a guide for rRNA/ribosome maturation enzymes to their substrates [14, 18].

### CshA and Era are both required for growth at suboptimal temperatures, virulence and rRNA processing

In *E. coli*, CshA has been linked to survival at low temperatures [34], while strains with depleted levels of Era are sensitive to both cold and heat shock [9, 10]. To examine the importance of these enzymes in *S. aureus*, we utilised deletion strains in the *S. aureus* MRSA strain LAC*. While both the *S. aureus era* and *cshA* null mutants showed significant growth defects at 37°C (Fig 1b and S3a), the growth of both was severely compromised at 25°C (Fig 4a and S3b). This defect was enhanced in a double Δ*era cshA* mutant, which failed to grow at 25°C even after 48 h (Fig 4a and S3b), highlighting that these two proteins are essential for bacteria to cope with growth at suboptimal temperatures.

**Fig 4.**
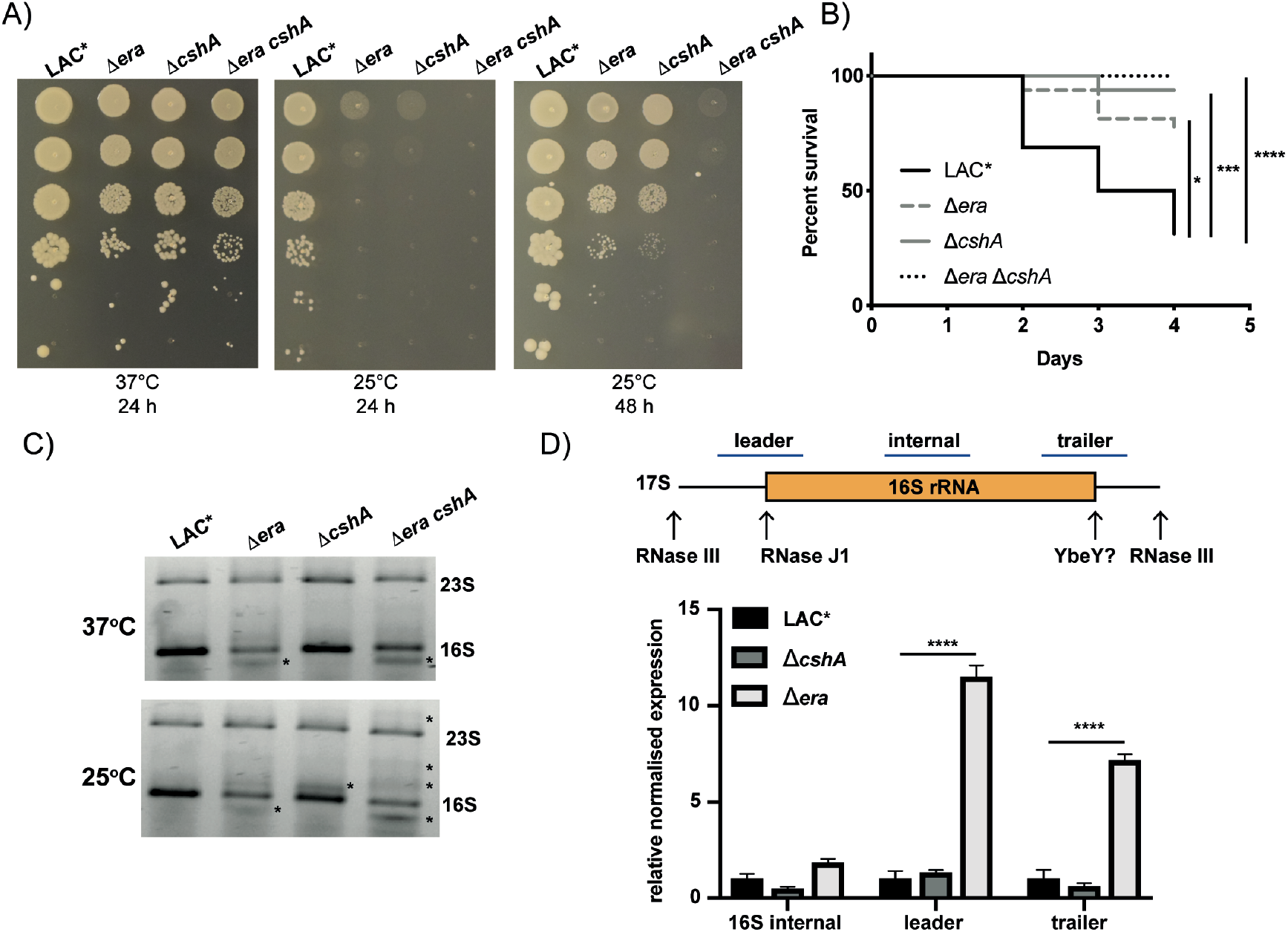
Era and CshA are required for cold shock survival, virulence and rRNA processing. A) Serial dilutions of the wildtype LAC*, Δ*era*, Δ*cshA* and Δ*era cshA* were spotted onto TSA agar plates and incubated at either 37 or 25°C for the times indicated. N=3 with one representative image shown. B) Survival of *G. mellonella* larvae inoculated into the hind leg with 1 × 10^5^ cfu of wildtype LAC* and mutant Δ*era*, Δ*cshA* and Δ*era cshA* strains. Larvae were incubated at 37°C for 4 days. Experiments were repeats in triplicate (N=3), with one graph shown. Survival curves were calculated by Kaplan-Meier analysis, with the log-rank (Mantel Cox) test performed to compare different strains (* *P* < 0.05, *** *P* < 0.0005, **** *P* < 0.0001). C) rRNA profiles from *era* and *cshA* mutant strains. 500 ng of RNA extracted from LAC*, Δ*era*, Δ*cshA* and Δ*era cshA* grown to an OD_600_ of 0.4 at either 37 or 25°C, were run on 0.7% agarose/0.9% synergel gels and stained with Evagreen dye. * highlights the presence of either processing or degradation intermediates. N=3 with one representative image shown. D) top: schematic of the 17S rRNA, with arrows indicating processing sites that lead to the formation of the mature 16S rRNA. The blue bars indicate the probe regions amplified by RT-PCR. Bottom: RT-PCR of total RNA from the wildtype LAC*, Δ*era*, and Δ*cshA* strains grown at 25°C using probes as indicated above. rRNA values are expressed as mean relative expression +/- SD normalised to the wildtype and the expression of *rho* RNA as an internal control. Statistical analysis was performed using a two-way ANOVA, followed by Tukey’s multiple comparisons test (**** *P* < 0.0001). Cycle threshold values were determined for 3 biological repeats in duplicate (N=3).

Era is an RA-GTPase that, along with a second RA-GTPase RsgA, binds to the 30S ribosomal subunit [14, 35]. In *E. coli*, overexpression of Era can complement a deletion of *rsgA* [36]. After confirming that expression of Era can complement a deletion of *rsgA* in *S. aureus* (Fig S4a), we used cross-complementation of the Δ*era* and Δ*cshA* mutations to investigate whether Era and CshA act in the same pathway and can functionally complement each other. However, cross-complementation did not improve growth at 25°C (Fig S4b and S4c), indicating that while these proteins interact and bacteria lacking both cannot survive cold temperatures, they have separate cellular functions. We next assessed the contribution of each gene to the virulence of *S. aureus* using *Galleria mellonella* as an invertebrate model of systemic infection. While the wildtype bacteria were fully virulent, both the *era* and *cshA* mutant strains displayed reduced killing, with the combined double mutant being completely avirulent (Fig 4b).

To understand why Era and CshA are so important for growth at cold temperatures and for virulence, we sought to assess the impact of deleting these genes on the cellular rRNA. We observed that RNA extracted from both the single Δ*era* and double Δ*era cshA* strains grown at 37°C had a processing or degradation intermediate migrating slightly below the 16S band (Fig 4c). At 25°C we also observed a defect for the Δ*cshA* mutant strain and multiple additional bands were evident in the Δ*era cshA* double mutant (Fig 4c), which could represent either processing intermediates from the 16S, or degradation intermediates from both the 23S and the 16S. A build-up of degradation products are expected in the *cshA* mutant due to its role in the RNA degradosome. However, to determine whether processing defects could be occurring in these mutant strains when grown at low temperatures, we performed qRT-PCR with probes that either amplified internally to the 16S rRNA sequence, or either flanked the 5’ or 3’ processing junctions required to convert 17S into mature 16S rRNA (Fig 4d). While the levels of 16S rRNA were not significantly different between wildtype and mutants, large increases in unprocessed 5’ and 3’ rRNA were evident in the Δ*era* strain (Fig 4d). As Era is known to interact with the 30S ribosomal subunit, this could potentially be due to mis-localisation of endonuclease YbeY in the absence of Era. It is therefore, possible that Era acts as an intermediary protein on the 30S ribosomal subunit, enabling proteins such as the endonuclease YbeY, and the RNA-degradosome helicase CshA to access the 17S rRNA, allowing for processing (maturation or degradation as appropriate) of rRNA under normal and stress conditions, such as growth at cold temperatures.

### Rel_SA_ interacts with both Era and CshA

Our previous work demonstrated that the GTPase activity of Era is inhibited by the stringent response alarmone ppGpp [22]. To examine whether ppGpp also interacts with CshA we performed DRaCALA assays, revealing that this nucleotide does not interact with CshA (Fig S5a), nor does the affinity of ppGpp for Era alter in the presence of CshA (K_d_ of 3.1 ± 0.4 μM in the absence versus 3.2 ± 0.3 μM in the presence of CshA) (Fig S5b).

To investigate whether the stringent response has any additional points of interaction with either Era and CshA, we examined whether one of the (p)ppGpp synthetases might directly interact with either protein via bacterial two-hybrid. This revealed that Rel_SA_, but not RelP or RelQ, interacts with CshA and that Rel_SA_ interacts with Era (Fig 5a). To confirm this interaction, we first used affinity pulldown assays, showing that although no significant interaction occurred with Era, CshA and Rel_SA_ interacted quite strongly (Fig 5b). Following this, we again used the split luciferase assay, which confirmed an interaction between Rel_SA_ and CshA (Fig 5c) or Era (Fig 5d). It was interesting to note that the CshA-Rel_SA_ interaction only occurred in the CshA_SB_-Rel_LB_ and not the Rel_SB_-CshA_LB_ orientation, highlighting the importance of creating both types of luciferase fusions.

**Fig 5.**
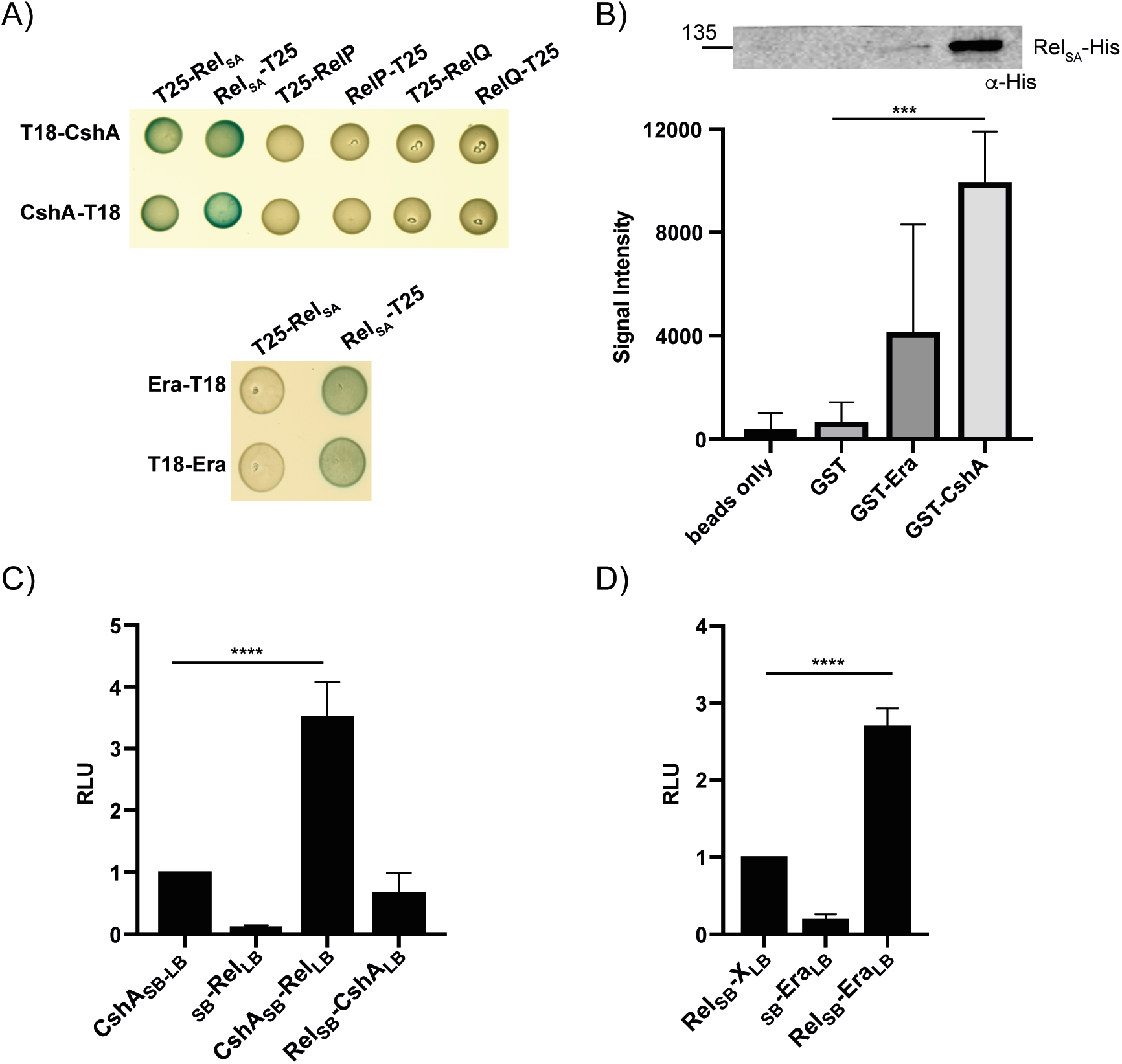
The stringent response synthetase Rel_SA_ interacts with Era and CshA. A) Top: Bacterial two-hybrid showing an interaction between CshA-T18/T18-CshA and both Rel_SA_-T25 and T25-Rel_SA_. CshA did not interact with the small (p)ppGpp synthetase enzymes RelP or RelQ. Bottom: Bacterial two-hybrid showing an interaction between Rel_SA_-T25 and both Era-T18 and T18-Era. N=3 with one representative image shown. B) Affinity pulldown assay using GST, GST-tagged Era and GST-tagged CshA coupled to glutathione beads. Beads alone, GST, GST-Era and GST-CshA coupled beads were incubated with His-tagged Rel_SA_. After washing, bound Rel_SA_-His was detected using HRP-conjugated anti-His antibodies. N=3 with one representative image shown above. Below, the mean signal intensities and standard deviations from the 3 repeats are plotted. Statistical analysis was performed using a one-way ANOVA, followed by Dunnett’s multiple comparisons test (*** *P* < 0.0005). C & D) Split luciferase assay demonstrating an interaction between CshA (C) or Era (D) and RelSA in *S. aureus*. The negative controls have genes singly fused to either the SB or the LB. The average values and standard deviations of triplicate experiments (N=3) are plotted. All statistical analysis was performed using a one-way ANOVA, followed by Dunnett’s multiple comparisons test (**** *P* < 0.0001).

### Rel_SA_ positively affects the GTPase activity of Era and is important for rRNA processing

We wished to ascertain whether the interactions between RelSA and Era impact the activity of either enzyme. Recombinantly purified full-length Rel_SA_ is known to be in the synthetase-off/hydrolase-on conformation *in vitro* [37]. Using purified Rel_SA_, we determined that Era and CshA have no effect on the pppGpp hydrolase activity of this enzyme (Fig S5c). However, upon examination of the GTPase activity of Era, we observed significantly increased activity in the presence of Rel_SA_ (Fig 6a).

**Fig 6.**
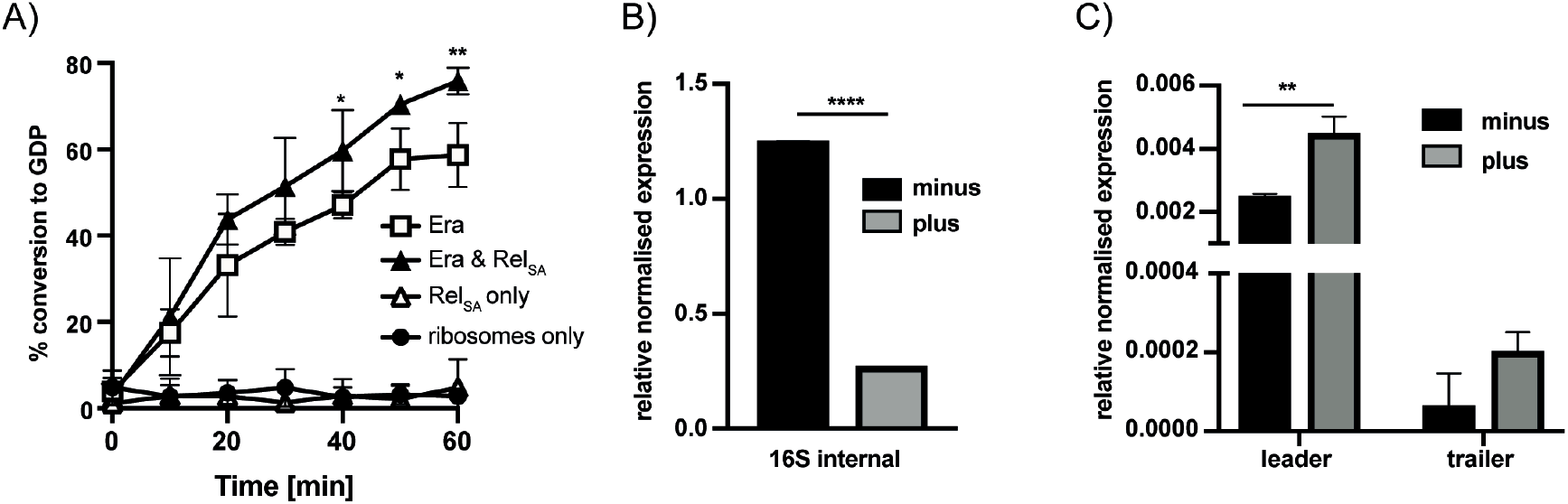
The stringent response influences Era GTPase activity and rRNA processing. A) The GTPase activity of 100 nM Era was measured in the presence of an equal amount of ribosomes, 100 nM Rel_SA_ and 1 μM GTP. All reactions contained ribosomes. The RelSA only control reaction contained RelSA and ribosomes but no Era. Hydrolysis of ^32^P-GTP was monitored by TLC and the percentage GDP formed quantified using ImageJ. Experiments were repeated three times with means and standard deviations shown. Statistical analysis was performed using a two-way ANOVA, followed by Dunnett’s multiple comparisons test (** *p* < 0.005, * *p* < 0.05). B & C) rRNA extracted from LAC*. Strains were grown at 25°C until an OD_600_ of 0.4. Cells were split and half exposed to 60 μg/ml mupirocin for 1 h (plus), which at 25°C corresponds to one doubling. RT-PCR of total RNA using the probes as indicated in Fig. 4d that bind internally to the 16S (B) or to the unprocessed leader and trailer sequences (C) of the 17S rRNA. rRNA values are expressed as mean relative expression +/- SD normalised to the expression of *rho* RNA as an internal control. For comparisons between two groups (B - LAC* minus v plus) a two-tailed, unpaired Student’s t-test was performed (**** *p* < 0.0001). For comparisons between more than two groups (C) a two-way ANOVA, followed by Sidak’s multiple comparisons test was performed (** *p* < 0.005). Cycle threshold values were determined for 3 biological repeats in duplicate (N=3).

Through Rel_SA_, the stringent response appears to interact with components of the ribosome and 16S rRNA processing machinery. To analyse the importance of the stringent response for rRNA processing, we grew the wildtype *S. aureus* LAC* at 25°C, a condition under which we have shown that both Era and CshA are important for viability, and induced the stringent response with mupirocin, an antibiotic that inhibits the isoleucyl-tRNA synthetase mimicking amino acid deprivation [38]. We again performed qRT-PCR with the probes as in Fig 4d that either amplified internally to the 16S rRNA sequence, or either flanked the 5’ or 3’ processing junctions of the 17S rRNA. For the probe internal to the 16S rRNA we observed a significant decrease in 16S rRNA in the presence of mupirocin (Fig 6b). This is expected, as it is known that rRNA levels decrease upon activation of the stringent response. However, when we compare the amount of unprocessed 5’ or 3’ sequences relative to the level of 16S in each strain, we observed a significant increase in the amount of unprocessed leader 17S rRNA upon induction of (p)ppGpp (Fig 6c: *p* = 0.0031). We have previously shown that ppGpp interacts with Era and that mature ribosome formation is inhibited upon induction of the stringent response [22]. Here we propose that the stringent response can also impact RA-GTPase function through direct interactions of RelSA with both Era and CshA, influencing the GTPase activity of Era and 17S rRNA processing.

## Discussion

The synthesis of ribosomes and proteins consumes approximately 40% of the energy within a growing bacterial cell [39]. Ribosomal assembly cofactors are, therefore, an essential group of enzymes for coordinating biogenesis as efficiently as possible. Members of this group include RNA helicases, rRNA and protein modification enzymes, chaperones and RA-GTPases. *S. aureus* contains over 11 RA-GTPases, each with potentially varying roles in the biogenesis of the 50S and 30S subunits, although the precise functions of each in this process is unclear.

While the RA-GTPase Era has been extensively studied, its precise function in the cell is unknown. Cryo-electron microscopy shows Era interacting with the 30S ribosomal subunit, however this protein has also been implicated in numerous other cellular processes [14]. Era-depleted *B. subtilis* or *E. coli* cells are elongated, with defects in septum formation. These cells also contain diffuse nucleoid material, implicating Era in cell division and chromosome segregation [5, 12]. In *B. subtilis*, Era-depleted cells have defects in spore formation [7], while growth defects in strains containing Era variants with reduced nucleotide binding abilities can be rescued by truncating either *rpoN* or *ptsN* [11]. RpoN is required for nitrogen assimilation and fixation, while PtsN is involved in sugar transport, suggesting that Era is also involved in regulating carbon and nitrogen metabolism. Localisation studies have indicated that this protein is present at both the membrane and in the cytoplasm [40], and so it has been suggested that this enzyme cycles between the membrane and the ribosome in response to cellular triggers. In addition to ribosomal biogenesis factors, Era has been reported to interact with MazG, Ndk, Pk and YggG [41–43]. Ndk, a nucleoside diphosphate kinase and Pk, a pyruvate kinase, from *Pseudomonas aeruginosa* were both shown to form a complex with Era, further implicating Era in energy metabolism [42]. MazG is a nucleoside triphosphate pyrophosphohydrolase, while YggG is a membrane-associated heat shock protein. The significance of these interactions with Era is unknown.

Era has also been implicated in helping cells cope with cold stress. Conditional cold-sensitive mutants of Era have been constructed in *E. coli* [10] and these mutations could be suppressed by the overexpression of the 16S rRNA methyltransferase KsgA, providing one of the first links between Era and ribosome biogenesis [9]. In addition, Era interacts with YbeY in *E. coli* [18]. YbeY is a universally conserved endonuclease that, while dispensable in *E. coli*, is essential in *B. subtilis* and potentially in *S. aureus*, given the lack of transposon mutants available [19, 25, 28]. This endonuclease is required for the maturation of the 3’ end of the 16S rRNA, and in strains depleted of YbeY the 70S ribosomes are targeted for degradation with the help of RNase R [19, 28]. Era and YbeY are fused in a polypeptide in clostridial species, further highlighting a link between these two proteins. Although not encoded in the same operon as *era* in *E. coli*, this prompted us to examine the interactions of all other proteins encoded in the *era* operon in *S. aureus* with Era (Fig 2). As in *E. coli*, YbeY from *S. aureus* interacts with Era. YbeZ also interacted by bacterial two-hybrid but not significantly using the quantitative luciferase assay (Fig 2). YbeZ has been shown to interact with YbeY in *E. coli* [18]. This protein has an ATPase domain and an RNA binding motif, and so is likely to also bind to the 17S rRNA and may form a complex with Era and YbeY to aid in processing the 17S rRNA to the mature 16S form.

Here we use interaction studies to show that Era interacts with a protein involved in survival at suboptimal temperatures, the DEAD-box RNA helicase CshA (Fig 3). In *B. subtilis*, CshA is one of the most abundant RNA helicases produced at low temperatures [44] and in *B. subtilis* and *E. coli* it has been implicated in multiple processes, including ribosome biogenesis and interacting with components of the RNA degradosome [32–34, 44]. In *S. aureus*, CshA has been linked to controlling the turnover of mRNA in the cell [31]. Deletion of *cshA* results in the stabilisation of some mRNA transcripts, such as the *spa* mRNA, but CshA also protects a number of other mRNA and sRNA transcripts under stress conditions [31]. In addition, the CT region of CshA is required for interactions with the degradosome [32]. Our analysis shows that Era and CshA interact *in vitro* and when expressed heterologously in *E. coli* (Fig 3a and 3b). We additionally used an adapted split luciferase assay to show this interaction natively in *S. aureus* (Fig 3c). Using this assay has also allowed us to determine that the CT disordered region of CshA, which is important for binding to mRNA and the degradosome [32], is dispensable for interacting with Era (Fig 3).

Using *cshA* and *era* deletion strains we show that both single mutants are important for growth at cold temperatures, with a double mutant unable to survive at 25°C (Fig 4). By examining the rRNA of these strains we observe that the *era* mutant strain has increased 17S unprocessed rRNA intermediates (Fig 4d). Additionally, a double *Dera cshA* mutant has multiple processing and degradation defects (Fig 4c). It has been proposed that Era may act to guide processing enzymes to their site of action [18]. In keeping with this, we have observed that these proteins have little effect on the enzymatic activity of each other. Therefore, we propose a model whereby Era acts as an intermediary protein, allowing proteins such as the endonuclease YbeY and the RNA helicase CshA, and potentially others, access to their substrate rRNA. It has been suggested that CshA helps RNase components of the degradosome access cleavage sites that may be otherwise inaccessible [45], and so it is tempting to speculate that Era is responsible for recruiting CshA to target misfolded ribosomal subunits for degradation under stress conditions.

Cellular levels of rRNA are controlled by the stringent response [46]. Upon activation of the stringent response, transcription from rRNA promoters is reduced (Fig 6b), either by direct binding of ppGpp to the RNA polymerase in Gram negatives, or by tight control of cellular GTP levels in Gram positives [47, 48]. Here we see an additional level of regulation by the stringent response on rRNA. We show a direct interaction between the (p)ppGpp synthetase Rel_SA_ and both CshA and Era (Fig 5), and show that RelSA has a positive effect on the GTPase activity of Era (Fig 6a). Additionally, we observed that the activation of the stringent response results in increased processing defects in a wildtype strain grown at 25°C (Fig 6c). RA-GTPases are reported to have increased association to the ribosome in the GTP-bound form. Hydrolysis of GTP then acts as a signal to promote dissociation, presumably following a maturation event [49, 50]. The increased GTPase activity of Era in the presence of RelSA could promote premature dissociation of Era from the ribosome, leading to processing defects (Fig 6c) and contributing to the stalled growth phenotype synonymous with the stringent response. The precise dynamics of processing-enzyme association under stress conditions requires further in-depth research but is beyond the scope of this study. For instance, under what growth conditions is Rel_SA_ binding to Era and CshA promoted and how does this affect the localisation of processing proteins to the ribosome?

Taken together, we demonstrate a cellular function for Era as a protein important for coordinating 30S ribosomal biogenesis in a stringent response-dependent manner. It is possible that defects in ribosomal assembly lead to altered protein translation, which may be the reason for the plethora of other phenotypes associated with depletion of Era. As Era can complement a defect in RsgA (Fig S4a, [36]), as well as RbfA [26], we propose a broader role for RA-GTPases as intermediary proteins. It is now of interest to determine what processing events are coordinated by the other RA-GTPases in *S. aureus*, as well as more broadly in other prokaryotes.

## Materials and methods

### Bacterial strains and culture conditions

*E. coli* strains were grown in Luria Bertani broth (LB) and *S. aureus* strains in tryptic soy broth (TSB) at 37°C with aeration. Strains and primers used are listed in Tables S1 and S2. Information on strain construction is provided in SI Methods.

### Luciferase assay

Overnight cultures of *S. aureus* strains were diluted to an OD_600_ of 0.05 and grown for 90 min in the presence of 100 ng/ml anhydrotetracycline (Atet). Strains were normalised to an OD_600_ of 0.1 and luciferase activity measured according to the Nano-Glo Luciferase Assay System Protocol (Promega).

### Pull-down experiments and western blotting

Glutathione beads were washed 5 times in 1 x wash solution (25 mM Tris pH 7.5, 150 mM NaCl, 1 mM EDTA, 0.5% Triton X-100). 1 μM of glutathione-S-transferase (GST), GST-CshA or GST-Era were coupled to the beads by incubating in 1 x wash buffer for 4 h at 4°C. Unbound protein was removed by washing in 1 x wash solution between 5-10 times. Protein-bound beads were incubated with 1 μM His-tagged Era, His-YbeY or His-Rel_SA_ in the presence of 1 x binding buffer (25 mM Tris pH 7.5, 150 mM NaCl, 5 mM MgCl_2_, 0.5% Triton X-100) at 4°C overnight. After washing, attached proteins were eluted from the beads with 50 μl elution buffer (25 mM Tris pH 8, 150 mM NaCl, 1 mM EDTA, 0.5% Triton X-100, 10 mM reduced glutathione). Samples were mixed 1:1 with 2 x SDS protein sample buffer. Aliquots were separated on 12% SDS-polyacrylamide gels and proteins subsequently transferred to PVDF membranes. Bound His-tagged proteins were detected using HRP-conjugated anti-His antibodies (Sigma) at a 1:500 dilution. Blots were developed by enhanced chemiluminescence and imaged using a ChemiDoc MP imager (Bio-Rad).

### Construction of bacterial two-hybrid library

Genomic DNA of *S. aureus* was extracted and partially digested by incubation with Sau3AI at 37°C for 20 min. Digested DNA was run on a 0.8% agarose gel and fragments of 500 to 1000 bp and 1000 to 3000 bp gel extracted and purified. This was repeated five times from separate genomic preps and DNA fragments pooled. The vector pUT18C was digested overnight at 37°C with BamHI and dephosphorylated with antarctic phosphatase for 90 min. Genomic DNA fragments were ligated into the pUT18C linearized vector using T4 ligase, transformed into *E. coli* DH5α competent cells (New England Biolabs) and plated onto LB agar with carbenicillin. Plates were scraped and transformant plasmids isolated using the GeneJET plasmid purification kit (Thermo scientific).

### Bacterial two-hybrid

Bacterial two hybrid plasmids containing genes of interest were co-transformed into *E. coli* BTH101 cells, plated on LB agar containing 150 μg/ml carbenicillin and 30 μg/ml kanamycin and incubated at 30°C overnight. Colonies were isolated, grown overnight at 30°C in LB broth containing 0.5 mM IPTG and 5 μl spotted onto LB agar containing 0.5 mM IPTG and 40 μg/ml X-gal. Plates were incubated at 30°C for up to 48 h. In some cases, BTH101 transformants were incubated overnight with 0.5 mM IPTG and spotted directly onto LB agar containing IPTG, X-gal and appropriate antibiotics.

### *Galleria mellonella* survival assays

*S. aureus* strains were grown overnight in TSB, pelleted at 17,000 x g, washed twice in sterile PBS and diluted to an OD_600_ of 0.1 (2-4 × 10^7^ cfu/ml). Colony numbers were confirmed by dilution and plating on TSA. Final larval stage *G. mellonella* weighing 250-350 mg (Live foods direct, Sheffield, UK) were stored in the dark at 4°C and used within 7 days. 5 μl of *S. aureus* inoculum was injected into the hemocoel through the last pro-leg using a Hamilton syringe 701 RN. Larvae were incubated in sealed plastic petri dishes at 37°C and the number of surviving individuals recorded daily.

### RNA extraction

Strains of *S. aureus* were grown overnight at 37°C and diluted to an OD_600_ of 0.05. Cultures were grown to an OD_600_ of 0.4 at either 37 or 25°C and harvested. RNA was extracted using the RiboPure RNA Purification Kit (Invitrogen) as per guidelines. RNA was visualised using a modified agarose gel containing 0.7% agarose and 0.9% Synergel (Diversified Biotech) in 0.5 X TBE (44.5 mM Tris, 1 mM EDTA pH 8, 44.5 mM boric acid) as per Wachi *et al*. 1999 [51]. RNA was visualised using EvaGreen fluorescent nucleic acid dye (Biotium) on a ChemiDoc MP imager (Bio-Rad).

### RT-PCR

RNA was extracted as described above. Complementary DNA was synthesised from 1.5 μg RNA with transcriptor reverse transcriptase (Sigma) and random primers. RT-PCR was performed on 100 ng of cDNA in triplicate using SYBR Green Jumpstart Taq readymix (Sigma). Primers RMC379/380 (16S internal), RMC570/571 (leader) and RMC572/548 (trailer) were designed to amplify 200-220 bp target regions in the 17S rRNA. The housekeeping gene *rho* was amplified with primers RMC573/574. The *rho* transcript was chosen as it has been shown to be highly stable under nutrient-deficient conditions under which the stringent response would be induced [52]. Cycle threshold values were determined for 3 biological repeats in duplicate. For each reaction, the ratio to *rho* transcript number was calculated as follows: 2^-(Ct target – Ct rho)^.

### Growth curves

*S. aureus* strains were grown overnight in TSB medium containing the appropriate antibiotics. Overnight cultures were diluted to a starting OD_600_ of 0.05 in the presence of 100 ng/ml Atet where appropriate, incubated at 25°C or 37°C with aeration and OD_600_ values determined at 2 h intervals. Growth curves were performed in triplicate, and averages and standard deviations are plotted.

### Minimum Inhibitory Concentrations

Overnight cultures of wildtype LAC* and Δ*era* strains were adjusted to an OD_600_ of 0.05 in Mueller-Hinton broth and 100 μl incubated in 96 well plates with 2-fold dilutions of various antimicrobials at the following starting concentrations: spectinomycin 250 μg/ml and chloramphenicol 64 μg/ml. Plates were incubated at 37°C overnight with shaking. Assays were performed in triplicate, and averages and standard deviations are plotted.

### Protein purifications

Proteins were purified from 1-2 L *E. coli* cultures. Cultures were grown to an OD_600_ 0.5-0.7, expression induced with 1 mM IPTG and incubated for 3 h at 37°C. Protein purifications were performed by either nickel or glutathione affinity chromatography. For nickel purifications, cell pellets were resuspended in 5 ml Buffer A (50 mM Tris pH 7.5, 150 mM NaCl, 5% glycerol, 10 mM imidazole), lysed with 100 μg lysozyme and sonication and the filtered cell lysate loaded onto a 1 ml HisTrap HP Ni^2+^ column (GE Healthcare) before elution using a gradient of Buffer B (50 mM Tris pH 7.5, 150 mM NaCl, 5% glycerol, 500 mM imidazole). GST-tagged proteins were resuspended in 5 ml PBS, lysed and the cell lysate was loaded onto a 1 ml GSTrap HP column (GE Healthcare) before elution using 50 mM Tris pH 8.0, 10 mM reduced glutathione. Protein containing fractions were dialysed in 50 mM Tris pH 7.5, 200 mM NaCl, 5% glycerol before storage at −80°C. Protein concentrations were determined by A_280_ readings.

### Synthesis of (p)ppGpp and differential radial capillary action of ligand assay (DRaCALA)

The synthesis of (p)ppGpp and DRaCALA binding assays were performed as described previously [22].

### Ribosomal profiles from *S. aureus* cell extracts

Crude isolations of ribosomes from *S. aureus* cell extracts were achieved as described by Chin Loh *et al*. with some modifications [27]. Briefly, 100 ml cultures of the different *S. aureus* strains were grown to an OD_600_ of 0.4 in TSB 100 ng/ml Atet. 100 μg/ml chloramphenicol was added to each culture, before being cooled to 4°C. Pelleted cells were suspended in association buffer (20 mM Tris-HCl pH 7.6, 8 mM MgCl_2_, 30 mM NH_4_Cl and 2 mM β-mercaptoethanol), normalized to an OD_600_ of 15, lysed by the addition of 0.2 μg/ml lysostaphin and 75 ng/ml DNase and incubated for 30 min at 37°C. The extracts were centrifuged at 17,000 x g for 5 min and subsequently 250 μl were layered onto 10-50% sucrose density gradients made in association buffer. Gradients were centrifuged for 4.5 h at 192,100 x g. Gradients were fractionated by upwards displacement of 250 μl aliquots, which were analysed for RNA content at an absorbance of 260 nm.

### Helicase assay

Reactions were set up as described [53]. Briefly, 250 nM of Cy3-labelled duplex RNA (Cy3-GCUUUACGGUGCUA, AACAAAACAAAAUAGCACCGUAAAGC) purchased from IDT, was incubated with 0.5 μM recombinant protein in reaction buffer (20 mM Tris pH 7.5, 50 mM potassium acetate, 5 mM magnesium acetate, 10 mM DTT, 0.1 mg/ml BSA, 10 U RNasin). The 0.5 μM recombinant protein used is at a concentration similar to that used in the affinity pulldown assays (1 μM) that show an Era-CshA interaction. The reaction was initiated with the addition of 1 mM ATP and incubated at 25°C for 10 min. Reactions were terminated by mixing with an equal volume of stop buffer (1% SDS, 0.5 mM EDTA, 20% glycerol) and loaded onto 20% native polyacrylamide gels before being electrophoresed in 0.5 x TBE. Bands were imaged using a ChemiDoc MP imager (Bio-Rad).

### Nucleotide hydrolysis assays

The ability of proteins to hydrolyse GTP was determined by incubating 100 nM recombinant Era with equimolar YbeY/CshA or Rel_SA_, 100 nM *S. aureus* ribosomes, 1 μM GTP and 2.78 nM a-^32^P-GTP in 40 mM Tris pH 7.5, 100 mM NaCl, 10 mM MgCl_2_ at 37°C for the indicated times. All reactions were set up in the absence of Era as controls. pppGpp hydrolysis assays were performed by incubating 100 nM recombinant Rel_SA_ in the presence and absence of equimolar GST, GST-Era or GST-CshA, 1 μM pppGpp and 2.78 nM a-^32^P-pppGpp in 50 mM Tris pH 8.5, 0.1 mM MnCl_2_, 20 mM KCl at 37°C for the indicated times. All reactions were set up in the absence of Rel_SA_ as controls. Hydrolysis reactions were inactivated with the addition of formic acid to a final concentration of 1.2 M. Precipitated proteins were pelleted by centrifugation at 17,000 x g for 10 min. Reaction products were then visualized by spotting 1 μl on PEI-cellulose thin layer chromatography (TLC) plates (Merck Millipore) and separated using 1 M KH_2_PO_4_, pH 3.6 buffer. The radioactive spots were visualised using an LA 7000 Typhoon PhosphorImager and images quantified using ImageJ.

## Supporting information

**S0 Fig. Antibiotic susceptibility testing**

**S1 Fig. Enzyme activity of CshA and Era**

**S2 Fig. Growth of Aera and Δ*cshA* strains at 37 and 25°C**

**S3 Fig. Cross-complementation of Δ*rsgA*, Δ*era* and Δ*cshA* strains**

**S4 Fig. ppGpp does not interact with CshA**

**Supplemental methods** – plasmid and strain construction

**S1 Table.** Bacterial strains used in this study

**S2 Table.** Primers used in this study

