## Supplemental figures, methods and tables for "The (p)ppGpp-binding GTPase Era promotes rRNA processing and cold shock survival in *Staphylococcus aureus*"

### Figures

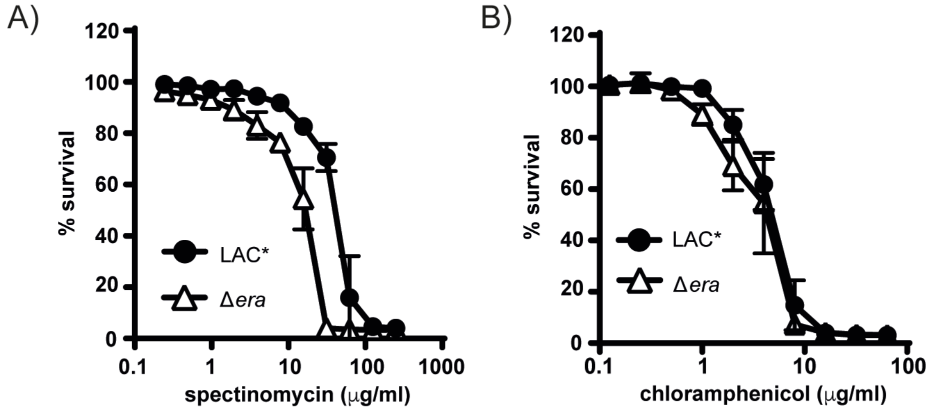


**S1 Fig. Antibiotic susceptibility testing.** The susceptibility of LAC* and Δ*era* to A) spectinomycin and B) chloramphenicol was measured by growing the strains in 96 well plates with the indicated concentration of each antibiotic. $OD_{600}$ readings were determined after 24 h of growth and plotted as % survival compared to growth without antibiotic. Experiments were repeated in triplicate and mean and standard deviation plotted.

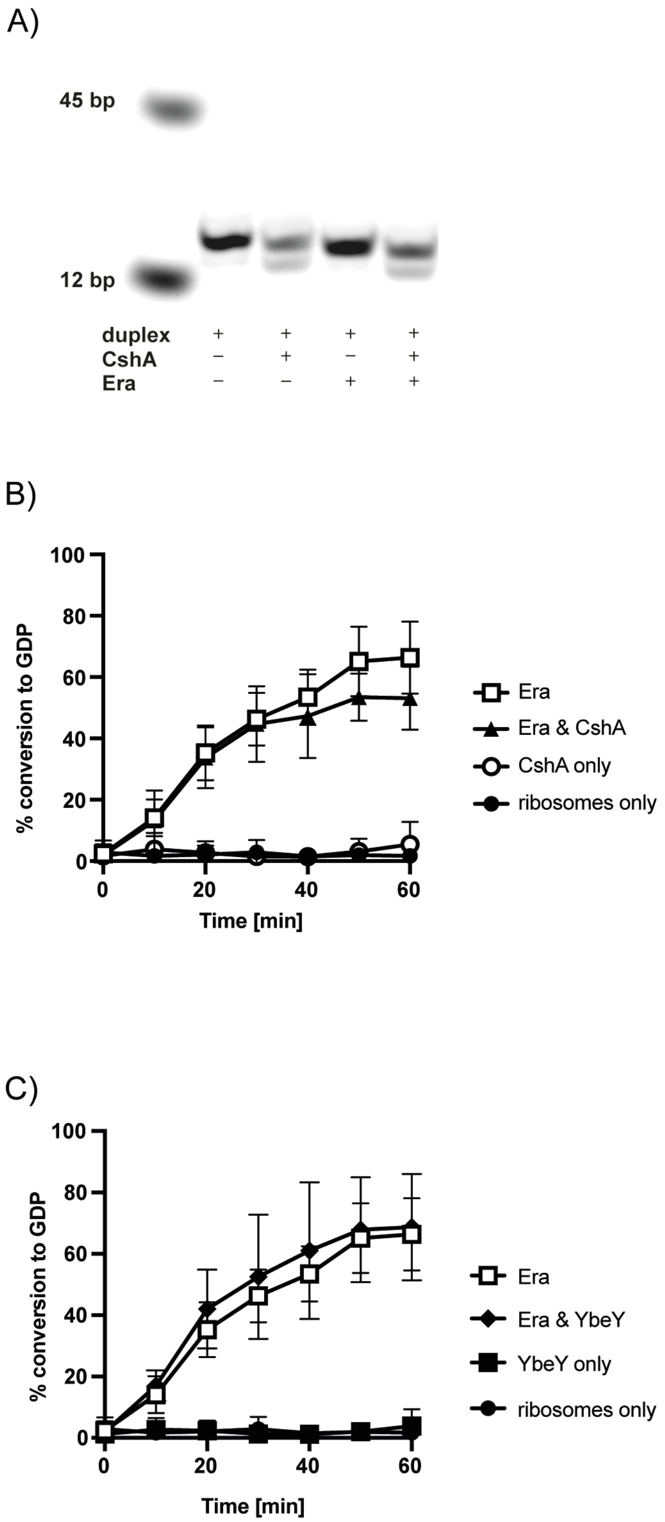


**S2 Fig. Enzyme activity of CshA and Era.** A) The RNA helicase activity of 0.5 μM CshA and Era were determined using a Cy3-labelled double stranded RNA oligomer. Reactions were incubated at 25°C for 10 min before analysis on a native page gel. B & C) The GTPase activity of 100 nM Era was measured in the presence of an equal amount of ribosomes and 1 μM GTP, plus and minus 100 nM CshA (B) or 100 nM YbeY (C). All reactions contained ribosomes. Reactions lacking Era were included as controls. Hydrolysis of $^{32}$P-GTP was monitored by TLC and the percentage GDP formed quantified using ImageJ. Experiments were repeated five times with means and standard deviations shown. Statistical analysis was performed using a two-way ANOVA, followed by Dunnett's multiple comparisons test.

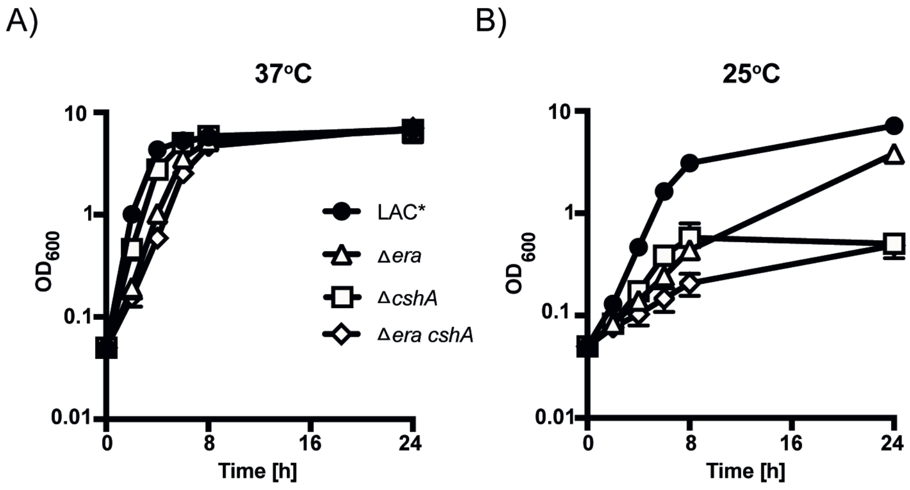


**S3 Fig. Growth of Δ*era* and Δ*cshA* strains at 37 and 25°C.** Growth of *S. aureus* strains LAC*, LAC* Δ*era,* LAC* Δ*cshA* and LAC* Δ*era cshA* at A) 37°C and B) 25°C. Overnight cultures were diluted to an $OD_{600}$ of 0.05 and grown for 24 h. Growth curves were performed in triplicate, with averages and standard deviations shown.

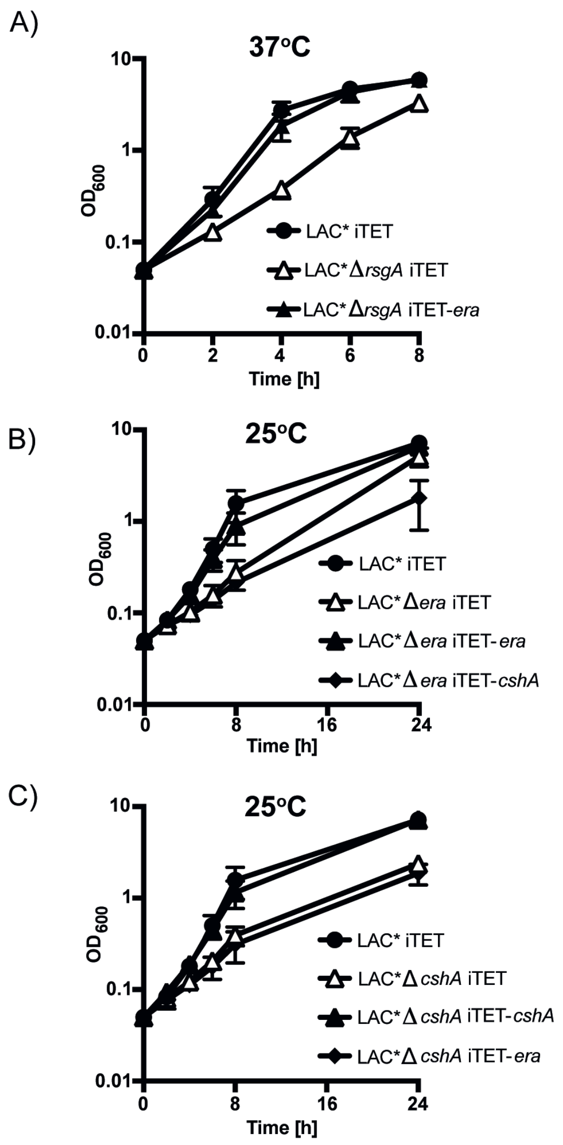


**S4 Fig. Cross-complementation of Δ*rsgA,* Δ*era* and Δ*cshA* strains.** A) Growth of *S. aureus* strains LAC* iTET, LAC* Δ*rsgA* iTET and LAC* Δ*rsgA* iTET-*era*. Overnight cultures were diluted to an $OD_{600}$ of 0.05 and grown in the presence of 100 ng/ml Atet for 8 h at 37°C. Era is able to complement the Δ*rsgA* growth defect. B) Growth of *S. aureus* strains LAC* iTET, LAC* Δ*era* iTET, LAC* Δ*era* iTET-*era* and LAC* Δ*era* iTET-*cshA*. Overnight cultures were diluted to an $OD_{600}$ of 0.05 and grown in the presence of 100 ng/ml Atet for 24 h at 25°C. C) Growth of *S. aureus* strains LAC* iTET, LAC* Δ*cshA* iTET, LAC* Δ*cshA* iTET-*cshA* and LAC* Δ*cshA* iTET-*era*. Overnight cultures were diluted to an $OD_{600}$ of 0.05 and grown in the presence of 100 ng/ml Atet for 24 h at 25°C. Growth curves were performed in triplicate, with averages and standard deviations shown.

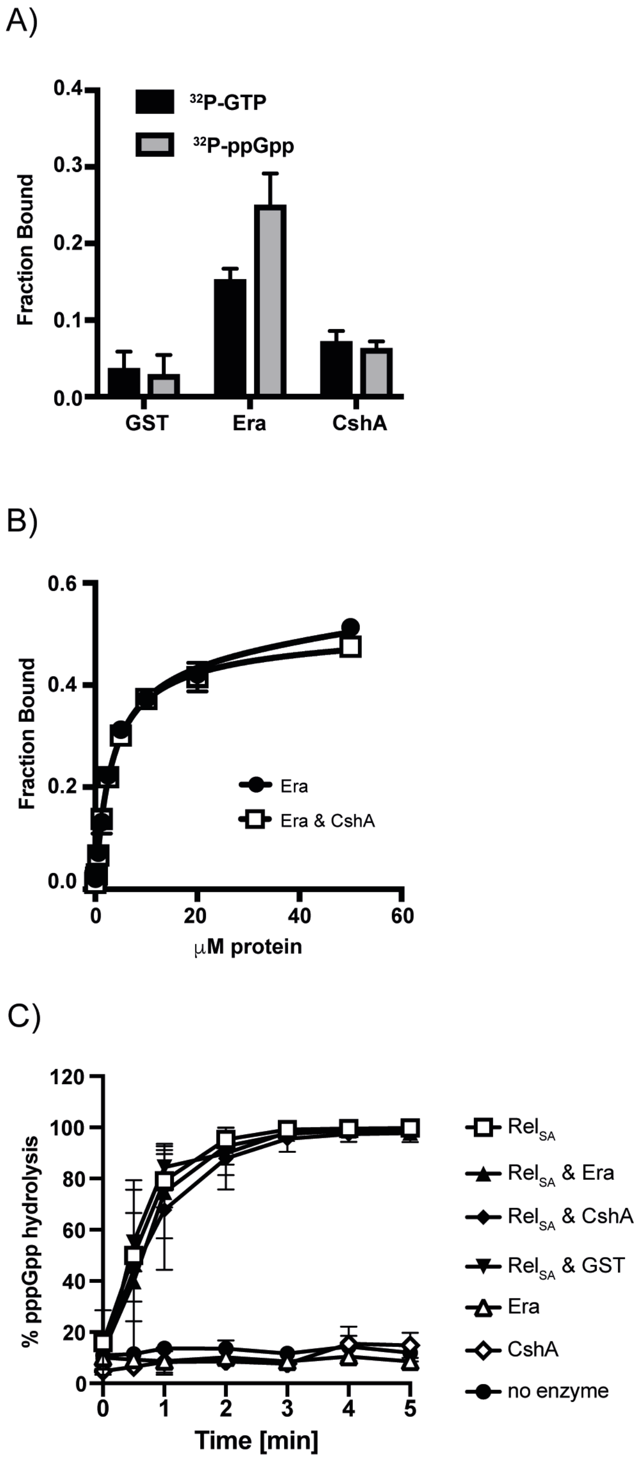


**S5 Fig. ppGpp does not interact with CshA.** A) DRaCALA binding assays with recombinant GST, GST-CshA and Era-His and $^{32}$P-labelled GTP and ppGpp. Quantification was carried out using ImageJ. The average values and standard deviations of triplicate experiments are plotted. B) Binding curves and $K_d$ determination for $^{32}$P-ppGpp and Era in the absence and presence of CshA. C) Hydrolysis activity of $Rel_{SA}$ on $^{32}$P-pppGpp in the absence and presence of Era and CshA. 100 nM of each protein were incubated with 1 μM pppGpp over the course of 5 min at 37°C before reactions were quenched. Reactions lacking $Rel_{SA}$, or including just GST in place of GST-Era/CshA were included as controls. Experiments were repeated in triplicate with means and standard deviations plotted. Statistical analysis was performed using a two-way ANOVA, followed by Dunnett's multiple comparisons test.

#### Supplemental Methods

**Plasmid and strain construction.** Strains used in this study are listed in Table S1 and primers are listed in Table S2. Strain construction is outlined below:

*S. aureus complementation plasmids*: Plasmid pCN55iTET was constructed by amplifying the iTET promoter from pRMC2 with primers RMC185/186 and cloning into the NarI/XmaI sites of pCN55. pCN55iTET-*era* and pCN55iTET-*cshA* were constructed by amplifying the *era* and *cshA* genes with primers RMC157/158 and RMC447/448 respectively, from LAC* genomic DNA and cloning into the KpnI/SacI sites of pCN55iTET.

*S. aureus gene deletion strains*: For the deletion of the *era* gene, 1 kb fragments up- and downstream of *era* were amplified from LAC* genomic DNA using primer pairs RMC178/151 and RMC154/179, which incorporate 5' and 3' BamHI sites. A *tetAM* gene was amplified from plasmid pTET using primer pair RMC152/153. Purified PCR products were then fused by SOE PCR using primers RMC178/179, digested with BamHI and cloned into the allelic exchange vector pTS1 yielding plasmid pTS1-Δ*era*. This plasmid was then electroporated into SEJ1 and stably maintained at 30°C in the presence of 10 μg/ml Cam. Shifting the temperature to 43°C resulted in insertion of the plasmid into the chromosome. The covering plasmid pCN55iTET-*era* was introduced into the strain, as it was at this point unknown if this gene was essential for growth or not. Downshift of the temperature to 30°C in the absence of chloramphenicol but in the presence of spectinomycin and Atet to select for and induce expression of Era from the plasmid, resulted in excision of the pTS1 plasmid and replaced the chromosomal copy of the *era* gene with the *tetAM* gene. The tetracycline marked *era* deletion was then transduced into a fresh LAC* strain background yielding LAC*Δ*era*.

*S. aureus luciferase strains*: plasmids pAP118-*era*$_{sb}$-*era*$_{lb}$, pAP118-*era*$_{sb}$-*cshA*$_{lb}$, pAP118-*era*$_{sb}$-*rel*$_{SAlb}$, pAP118-*rel*$_{SAsb}$-*era*$_{lb}$, pAP118-*rel*$_{SAsb}$-*cshA*$_{lb}$, pAP118-*cshA*$_{sb}$-*era*$_{lb}$, pAP118-*cshA*$_{sb}$-*rel*$_{SAlb}$, pAP118-*era*$_{sb}$-*cshA* 1-221$_{lb}$, pAP118-*era*$_{sb}$-*cshA* 1-382$_{lb}$, pAP118-*era*$_{sb}$-*cshA* 222-382$_{lb}$, pAP118-*era*$_{sb}$-*cshA* 383-506$_{lb}$, pAP118-*era*$_{sb}$-*ybeZ*$_{lb}$, pAP118-*era*$_{sb}$-*ybeY*$_{lb}$, pAP118-*era*$_{sb}$-*dgkA*$_{lb}$, pAP118-*era*$_{sb}$-*cdd*$_{lb}$, pAP118-*era*$_{sb}$-*recO*$_{lb}$, and pAP118-*era* 1-180$_{sb}$-*cshA*$_{lb}$ were constructed by first amplifying either the full-length *era, rel*$_{SA}$ or *cshA* genes, or the *era* 1-180 amino acid fragment, with the primers listed in Table S2 and cloning into the SacI/XhoI sites of pAP118, fusing each gene to the small bit (*sb*) of the nanoluc luciferase gene. pAP118 vectors were subsequently digested with PvuI and NotI. The appropriate genes for fusing to the large bit (*lb*) of the nanoluc were amplified with primers as specified in Table S2 and cloned into digested pAP118 vectors. The control strains pAP256-*era*$_{sb}$, pAP256-*rel*$_{SAsb}$, pAP256-*cshA*$_{sb}$, pAP256-*era* 1-180$_{sb}$ and pAP257-*cshA*$_{lb}$ were created by amplifying the respective genes and cloning into pAP256 or pAP257. All plasmids were electroporated into RN4220 Δ*spa,* before isolation and electroporation into the appropriate background.

*E. coli strains and plasmids:* All pKT25, pKNT25, pUT18 and pUT18C plasmids were constructed by amplifying the appropriate gene from LAC* genomic DNA with the primers listed in Table S2. PCR products were digested with XbaI/KpnI and cloned into digested vector. Plasmid pET28b-*era* for expression of His-Era was constructed by amplifying *era* with primers RMC68/69 and cloning into the BamHI and NdeI sites of digested pET28b. The GST-CshA and GST-Era producing clones pGEX-2TK-*cshA* and pGEX-2TK-*era* were created by cloning full-length *cshA* and *era* with the primers RMC401/402 and RMC657/658 into the SmaI site for *cshA* and BamHI/EcoRI sites for *era* of pGEX-2TK, respectively. All plasmids were initially transformed into *E. coli* strain XL1-Blue and sequences of all inserts were verified by fluorescence automated sequencing by GATC. For protein expression and purification, all pET28b and pGEX-2TK derived plasmids were transformed into *E. coli* strain BL21 (DE3).

#### S1 Table. Bacterial strains used in this study

| Strain | Relevant features | Reference |
|---|---|---|
| | ***Escherichia coli* strains** | |
| XL1-Blue | Cloning strain: TetR | Stratagene |
| DH5α | Cloning strain | [1] |
| BL21 (DE3) | Strain used for protein expression | Novagen |
| BTH101 | BACTH Δ*cya* strain | Euromedex |
| RMC0101 | pTS1 in DH5α | [2] |
| RMC0109 | pCN55 in XL1-Blue: CarbR | [3] |
| RMC0114 | pGEX-2TK in DH5α: CarbR | Pharmacia |
| RMC0127 | pKT25 in XL1 Blue: KanR | [4] |
| RMC0128 | pKNT25 in XL1 Blue: KanR | [5] |
| RMC0129 | pUT18 in XL1 Blue: CarbR | [4] |
| RMC0130 | pUT18C in XL1 Blue: CarbR | [4] |
| RMC0138 | pRMC2 in XL1 Blue: CarbR | [6] |
| RMC0147 | pET28b in XL1-Blue: KanR | Novagen |
| RMC0172 | pVL847-$rel_{sa}$ in T7IQ: GnR, CamR | [7] |
| RMC1453 | pVL791-*ybeY* in T7IQ: CarbR, CamR | [7] |
| RMC0397 | pET28b-*era* in XL1-Blue: KanR | This study |
| RMC0482 | pTS1-Δ*era* in XL1-Blue: CarbR | This study |
| RMC0531 | pCN55iTET in XL1-Blue: iTET promoter in pCN55; CarbR | This study |
| RMC0533 | pCN55iTET-*era* in XL1-Blue: CarbR | This study |
| RMC0935 | pCN55iTET-*cshA* in XL1-Blue: CarbR | This study |
| RMC0693 | pKT25-$rel_{sa}$ in DH5α: T25 fused to the N-terminus of $rel_{sa}$: KanR | This study |
| RMC0694 | pKNT25-$rel_{sa}$ in DH5α: T25 fused to the C-terminus of $rel_{sa}$: KanR | This study |
| RMC0697 | pKT25-*relP* in DH5α: T25 fused to the N-terminus of *relP*: KanR | This study |
| RMC0698 | pKNT25-*relP* in DH5α: T25 fused to the C-terminus of *relP*: KanR | This study |
| RMC0701 | pKT25-*relQ* in DH5α: T25 fused to the N-terminus of *relQ*: KanR | This study |
| RMC0702 | pKNT25-*relQ* in DH5α: T25 fused to the C-terminus of *relQ*: KanR | This study |
| RMC0744 | pKNT25-HD aa41-187 of $Rel_{SA}$ in DH5α: KanR | This study |
| RMC0748 | pKNT25-SYN aa247-357 of $Rel_{SA}$ in DH5α: KanR | This study |
| RMC0752 | pKNT25-TGS aa402-461 of $Rel_{SA}$ in DH5α: KanR | This study |
| RMC0756 | pKNT25-ACT aa657-735 of $Rel_{SA}$ in DH5α: KanR | This study |
| RMC0817 | pUT18C-*era* in BTH101: CarbR | This study |
| RMC0821 | pUT18-*era* in BTH101: CarbR | This study |
| RMC0825 | pKNT25-*era* in BTH101: KanR | This study |
| RMC0829 | pKT25-*era* in BTH101: KanR | This study |
| RMC0910 | pUT18C-*cshA* in XL1-Blue: T18 fused to the N-terminus of *cshA*: CarbR | This study |

| | | |
|---|---|---|
| RMC0911 | pUT18-*cshA* in XL1-Blue: T18 fused to the C-terminus of *cshA*: CarbR | This study |
| RMC1260 | pUT18C-*ybeZ* in XL1-Blue: T18 fused to the N-terminus of *ybeZ;* CarbR | This study |
| RMC1201 | pUT18C-*ybeY* in XL1-Blue: T18 fused to the N-terminus of *ybeY*: CarbR | This study |
| RMC1202 | pUT18C-*dgkA* in XL1-Blue: T18 fused to the N-terminus of *dgkA*: CarbR | This study |
| RMC1203 | pUT18C-*recO* in XL1-Blue: T18 fused to the N-terminus of *recO*: CarbR | This study |
| RMC1204 | pUT18C-*cdd* in XL1-Blue: T18 fused to the N-terminus of *cdd*: CarbR | This study |
| RMC0909 | pGEX-2TK–*cshA* in XL1-Blue: CarbR | This study |
| RMC1445 | pGEX-2TK–*era* in XL1-Blue: CarbR | This study |
| RMC0945 | pAP118 in DH5α: CamR | [8] |
| RMC0946 | pAP256 in DH5α: CamR | [8] |
| RMC0947 | pAP257 in DH5α: CamR | [8] |
| RMC1098 | pAP118-$era_{sb}$-$era_{lb}$ in XL1-Blue: CamR | This study |
| RMC1099 | pAP118-$era_{sb}$-$cshA_{lb}$ in XL1-Blue: CamR | This study |
| RMC1102 | pAP257-$cshA_{lb}$ in XL1-Blue: CamR | This study |
| RMC1156 | pAP118-$era_{sb}$-*cshA* $383\text{-}506_{lb}$ in XL1-Blue: CamR | This study |
| RMC1157 | pAP118-$era_{sb}$-*cshA* $1\text{-}382_{lb}$ in XL1-Blue: CamR | This study |
| RMC1158 | pAP257-*cshA* $383\text{-}506_{lb}$ in XL1-Blue: CamR | This study |
| RMC1159 | pAP257-*cshA* $1\text{-}382_{lb}$ in XL1-Blue: CamR | This study |
| RMC1160 | pAP118-$era_{sb}$-$ybeZ_{lb}$ in XL1-Blue: CamR | This study |
| RMC1161 | pAP118-$era_{sb}$-$ybeY_{lb}$ in XL1-Blue: CamR | This study |
| RMC1162 | pAP118-$era_{sb}$-$dgkA_{lb}$ in XL1-Blue: CamR | This study |
| RMC1163 | pAP118-$era_{sb}$-$cdd_{lb}$ in XL1-Blue: CamR | This study |
| RMC1164 | pAP118-$era_{sb}$-$recO_{lb}$ in XL1-Blue: CamR | This study |
| RMC1221 | pAP118-*era* $1\text{-}180_{sb}$- $cshA_{lb}$ in XL1-Blue: CamR | This study |
| RMC1222 | pAP256-*era* $1\text{-}180_{sb}$ in XL1-Blue: CamR | This study |
| RMC1236 | pAP257-*cshA* $1\text{-}221_{lb}$ in XL1-Blue: CamR | This study |
| RMC1244 | pAP118-$era_{sb}$-*cshA* $1\text{-}221_{lb}$ in XL1-Blue: CamR | This study |
| RMC1272 | pAP257-*cshA* $222\text{-}382_{lb}$ in XL1-Blue: CamR | This study |
| RMC1287 | pAP118-$era_{sb}$-*cshA* $222\text{-}382_{lb}$ in XL1-Blue: CamR | This study |
| RMC1444 | pAP118-$cshA_{sb}$-$era_{lb}$ in XL1-Blue: CamR | This study |
| RMC1443 | pAP118-$rel_{SAsb}$-$era_{lb}$ in XL1-Blue: CamR | This study |
| RMC1454 | pAP118-$era_{sb}$-$rel_{SAlb}$ in XL1-Blue: CamR | This study |
| RMC1455 | pAP118-$cshA_{sb}$-$rel_{SAlb}$ in XL1-Blue: CamR | This study |
| RMC1143 | pAP118-$rel_{SAsb}$-$cshA_{lb}$ in XL1-Blue: CamR | This study |
| RMC1456 | pAP256-$cshA_{sb}$ in XL1-Blue: CamR | This study |
| RMC1138 | pAP256-$rel_{SAsb}$ in XL1-Blue: CamR | This study |
| RMC1259 | pAP256-$era_{sb}$ in XL1-Blue: CamR | This study |

| | | |
|---|---|---|
| | ***Staphylococcus aureus* strains** | |
| SEJ1 | RN4220 Δ*spa;* protein A negative derivative of RN4220 | [9] |
| LAC* | Erm sensitive CA-MRSA LAC strain | [10] |
| NE565 | USA300 strain JE2 with transposon insertion in *cshA*: ErmR | [11] |
| RMC0358 | LAC* Δ*rsgA*: ErmR | [12] |
| RMC0447 | RN4220 pTET: TetR | [10] |
| RMC0552 | RN4220Δ*spa* Δ*era* pCN55iTET-*era*: TetR, SpecR | This study |
| RMC0562 | LAC* pCN55iTET: SpecR | This study |
| RMC0650 | LAC* Δ*era*: TetR | This study |
| RMC0813 | LAC* Δ*era* pCN55iTET: TetR, SpecR | This study |
| RMC0814 | LAC* Δ*era* pCN55iTET-*era*: TetR, SpecR | This study |
| RMC0908 | LAC* Δ*cshA*: ErmR | This study |
| RMC1030 | LAC* Δ*cshA* pCN55iTET: ErmR, SpecR | This study |
| RMC1031 | LAC* Δ*cshA* pCN55iTET-*era*: ErmR, SpecR | This study |
| RMC1032 | LAC* Δ*cshA* pCN55iTET-*cshA*: ErmR, SpecR | This study |
| RMC1033 | LAC* Δ*cshA* Δ*era*: ErmR TetR | This study |
| RMC1034 | LAC* Δ*era* pCN55iTET-*cshA*: TetR, SpecR | This study |
| RMC0890 | LAC* Δ*rsgA* pCN55iTET: ErmR, SpecR | This study |
| RMC0891 | LAC* Δ*rsgA* pCN55iTET-*era*: ErmR, SpecR | This study |

Antibiotics were used at the following concentrations - for *E. coli* cultures: carbenicillin (CarbR) 50-150 μg/ml; kanamycin (KanR) 30 μg/ml; chloramphenicol (CamR) 20 μg/ml, gentamicin (GnR) 20 μg/ml. For *S. aureus* cultures: erythromycin (ErmR) 10 μg/ml; chloramphenicol (CamR) 7.5 to 10 μg/ml; spectinomycin (SpecR) 250 μg/ml; tetracycline (TetR) 2 μg/ml. IPTG was used at 1 mM and anhydrotetracycline (Atet) at 100 ng/ml.

#### S2 Table. Primers used in this study

| Number | Name | Sequence |
|---|---|---|
| RMC068 | F-NdeI-Era | CCCCATATGACAGAACATAAATCAGCATTTGT |
| RMC069 | R-BamHI-Era | CCCGGATCCTTAATCTTGGTCTTCAACATAACC |
| RMC178 | F-BamHI-Era | AAAGGATCCGAAGAAGATGAGCCAGAGATTG |
| RMC179 | R-BamHI-Era | AAAGGATCCCACCTTATTGTGTAACATTCGAC |
| RMC151 | R-up FtetAM | AATAATTTTCATTTATTCTAAATCCTTTCCTGAAAA |
| RMC152 | F-tetAM | GATTTAGAATAAATGAAAATTATTAATATTGGAGTT |
| RMC153 | R-tetAM | CCACTTTTAAGACTAAGTTATTTTATTGAACATATA |
| RMC154 | F-down Rtet | AAAATAACTTAGTCTTAAAAGTGGTGAAGATAATTG |
| RMC157 | F-KpnI-Era | GGGGGGTACCTCAGGAAAGGATTTAGAATAAATG |
| RMC158 | R-SacI-Era | CCCCGAGCTCTTAATCTTGGTCTTCAACATAACC |
| RMC185 | R-NarI-tetR | CCTTGGCGCCTTAAGACCCACTTTCACATTTAAGTTG |
| RMC186 | F-XmaI-pRMC2 | TCCCCCGGGCGGAATTCGAGCTCAGATCTGTTAACGGTACCATC |
| RMC258 | F-XbaI-$Rel_{SA}$ | GGGGTCTAGAGAACAACGAATATCC |
| RMC259 | R-KpnI- $Rel_{SA}$ | GGGGGTACCTTCCAAACTCTTGTTAC |
| RMC260 | F-XbaI-RelP | GGGTCTAGAGTATGTAGATCGAAAACC |
| RMC261 | R-KpnI-RelP | GGGGGTACCTCTGTTATTTCAGAATG |
| RMC262 | F-XbaI-RelQ | GGGTCTAGAGAATCAATGGGATCAG |
| RMC263 | R-KpnI-RelQ | GGGGGTACCTCATTTTCATGTTTTTTAGAACG |
| RMC266 | F-XbaI-HD | GGGTCTAGATATTGCTTATGAAGCAC |
| RMC267 | R-KpnI-HD | GGGGGTACCTTAATACCAAGACGATGTG |
| RMC268 | F-XbaI-Synth | GGGTCTAGATGGTAGACCTAAACATATTTAC |
| RMC269 | R-KpnI-Synth | GGGGGTACCTTTTTACCTTCTTTGTAAGC |
| RMC270 | F-XbaI-TGS | GGGTCTAGAAGTATACGCATTTACCCC |
| RMC271 | R-KpnI-TGS | GGGGGTACCCTAGTACGTATTTCAAC |
| RMC272 | F-XbaI-ACT | GGGTCTAGATCAAAAATATCAGGTTG |
| RMC326 | R-KpnI-Era | CCCGGTACCCCATCTTGGTCTTCAACATAACCAATTTG |
| RMC330 | F-XbaI-Era | GGGTCTAGAGACAGAACATAAATCAGGATTTGTTTC |
| RMC401 | F-SmaI-CshA | AGTCCCCGGGGCAAAATTTTAAAGAACTAG |
| RMC402 | R-SmaI-CshA | AGCTCCCGGGTTTTTGATGGTCAGCAAATGTG |
| RMC407 | F-XbaI-CshA | GGGTCTAGATTGCAAAATTTTAAAGAACTAGGGATTTC |
| RMC408 | R-KpnI-CshA | CCCGGTACCCCTTTTTGATGGTCAGCAAATGTGC |
| RMC434 | F-XbaI-YbeZ | GGGTCTAGAGATGAAAAGGAGCGCGTG |
| RMC435 | R-KpnI-YbeZ | CCCGGTACCGGATTCTCTCCTTCATAATGTTCAATGATC |
| RMC436 | F-XbaI-YbeY | GGGTCTAGAGATGTTTACGATAGATTTTAGCGATC |
| RMC437 | R-KpnI-YbeY | CCCGGTACCGGGTCTCGTGTTAATCCATATGCG |
| RMC438 | F-XbaI-DgkA | GGGTCTAGAGATGAAAAGGTTTAAATATGCACTTG |

| | | |
|---|---|---|
| RMC439 | R-KpnI-DgkA | CCCGGTACCGGAAATAACGCTATAAAATGTGGTAAAAATAC |
| RMC440 | F-XbaI-RecO | GGGTCTAGAGATGCGCCAAAAAGGGATTATC |
| RMC441 | R-KpnI-RecO | CCCGGTACCGGTTGTTCCAATCTTTTTAATTGGTTG |
| RMC442 | F-XbaI-Cdd | GGGTCTAGAGATGAGTTATCAACCTCATTATTTTCAAG |
| RMC443 | R-KpnI-Cdd | CCCGAATCCGGTTCTAAATCCTTTCCTGAAAATCC |
| RMC447 | F-KpnI-CshA | GGGGGGTACCCAGGTAAAAAGGAGAATTATTTTG |
| RMC448 | R-SacI-CshA | CCCCGAGCTCTTATTTTTGATGGTCAGCAAATGT |
| RMC483 | F-SacI-Era | GGGGAGCTCTTTCAGGAAAGGATTTAGAATAAA |
| RMC484 | R-XhoI-Era | GGGCTCGAGAATCTTGGTCTTCAACATAACCAA |
| RMC485 | F-PvuI-Era | GGGCGATCGTTTCAGGAAAGGATTTAGAATAAA |
| RMC486 | R-NotI-Era | GGGGCGGCCGCATCTTGGTCTTCAACATAACCAAT |
| RMC487 | F-PvuI-CshA | GGGCGATCGCAGGTAAAAAGGAGAATTATTTTG |
| RMC488 | R-NotI-CshA | GGGGCGGCCGCTTTTTGATGGTCAGCAAATGTGC |
| RMC497 | R-NotI-CshA382 | GGGGCGGCCGCGATGTCATCTTCACGTGCTTG |
| RMC498 | F-PvuI-CshA383 | GGGCGATCGAAAAAGGAGAATTATTAAAGAAAAAGTTGAAAACTGGATG |
| RMC499 | F-PvuI-YbeZ | GGGCGATCGGCATCATAGAATGAATATAAATGATAT |
| RMC500 | R-NotI-YbeZ | GGGGCGGCCGCATTCTCTCCTTCATAATGTTCAATG |
| RMC501 | F-PvuI-YbeY | GGGCGATCGGAACATTATGAAGGAGAGAATTAA |
| RMC502 | R-NotI-YbeY | GGGGCGGCCGCGTCTCGTGTTAATCCATATGC |
| RMC503 | F-PvuI-DgkA | GGGCGATCGCATATGGATTAACACGAGACTAATT |
| RMC504 | R-NotI-DgkA | GGGGCGGCCGCAAATAACGCTATAAAATGTGG |
| RMC505 | F-PvuI-Cdd | GGGCGATCGCGTTATTTTAGGGAGGCATAT |
| RMC506 | R-NotI-Cdd | GGGGCGGCCGCTTCTAAATCCTTTCCTGAAAATCC |
| RMC507 | F-PvuI-RecO | GGGCGATCGCTTAAAAGTGGTGAAGATAATTGTTA |
| RMC508 | R-NotI-RecO | GGGGCGGCCGCTTGTTCCAATCTTTTTAATTGGTTG |
| RMC516 | R-XhoI-Era 1-180 | GGGCTCGAGATCATCTGGATAATATTTAGGTCC |
| RMC517 | R-NotI-CshA221 | GGGGCGGCCGCAATTGTATAGAATTCTTCGATTTG |
| RMC533 | F-PvuI-CshA222 | GGGCGATCGAAAAAGGAGAATTATTGTTAAAGAATTAGAGAAATTTGATAC |
| RMC570 | 5'-Leader-F | TTAGTATTTATGAGCTAATCAAACATC |
| RMC571 | 5'-Leader-R | AAAATATTATCCGGTATTAGCTCC |
| RMC379 | Internal-F | AGCTTAGTTGCCATCATTAAGTTGG |
| RMC380 | Internal-R | GTTGCAGACTACAATCCGAACTG |
| RMC572 | 3'trailer-F | CATGCTACGGTGAATACGTT |
| RMC548 | 3'trailer-R | ACGTTATTCCGCATCTTCTG |
| RMC573 | Rho-F | GAAGCTGCTGAAGTCG |
| RMC574 | Rho-R | GAATGCTTTTGGTTTGTGTAA |
| RMC657 | F-BamHI-Era | GGGGGATCCATGACAGAACATAAATCAGGATTTG |
| RMC658 | R-EcoRI-Era | GGGGAATTCTTAATCTTGGTCTTCAACATAACC |
| RMC659 | R-XhoI-CshA | GGGCTCGAGATTTTTGATGGTCAGCAAATGTGC |

Restriction sites in primer sequences are underlined
